# Peripheral nerve injury reallocates primary afferent input through spinal parvalbumin microcircuits

**DOI:** 10.64898/2026.07.15.737290

**Authors:** Haoyi Qiu, Hendrik Wildner, Hanns Ulrich Zeilhofer, Radhouane Dallel, Cedric Peirs, Reza Sharif-Naeini

## Abstract

Complex neural functions rely on finely tuned circuits in which the recruitment of local interneurons gates the flow of information, determining whether an input is relayed, amplified, or suppressed. Somatosensory information, such as touch or pain, is processed through such complex circuits in the dorsal horn of the spinal cord. There, local inhibitory interneurons play a key role in the proper segregation of touch and pain inputs. After nerve injury-evoked neuropathic pain, loss of inhibition impairs the function of these circuits, resulting in mechanical allodynia, where innocuous touch is perceived as painful. Disinhibition can be attributed to pruning of inhibitory synapses, reduced intrinsic excitability of inhibitory neurons, or weakened excitatory drive from primary afferents. Yet the complexity of the excitatory drive onto inhibitory neurons, and its potential modification after nerve injury, remains largely unexplored. Here we examined the nature of the synaptic drive from low-threshold Aβ mechanoreceptors (Aβ-LTMRs) onto parvalbumin-expressing interneurons (PVNs), and how this recruitment is affected by nerve injury. Aβ-LTMR stimulation evoked excitatory responses in a subset of PVNs, which is expected given the exclusively excitatory nature of primary afferent inputs. However, an unexpected subset of PVNs displayed inhibitory responses, suggesting the recruitment of a feedforward inhibitory circuit. We reconciled these observations by showing that Aβ-LTMRs engaged PVNs through both direct excitation and feedforward inhibition, which are differentially distributed between the inhibitory (iPVN) and excitatory (ePVN) subpopulation. Indeed, our results show that under naïve conditions, Aβ input preferentially recruited iPVNs, while ePVNs were predominantly suppressed by Aβ-driven feedforward inhibition mediated by a previously unrecognized Complexin-1 (Cplx1)-expressing inhibitory interneuron. After peripheral nerve injury, this balance becomes functionally redistributed. Aβ-to-iPVN transmission showed increased failure rates and impaired temporal precision, whereas Aβ drive onto ePVNs shifted from inhibition toward excitation. Notably, these functional changes occurred despite preserved afferent connectivity, synapse density, and spontaneous synaptic events, indicating that the dynamic reallocation of circuit recruitment occurs in the absence of structural changes. To test the behavioral consequences of these two populations, we used chemogenetic approaches and found that iPVNs suppress, whereas ePVNs promote, mechanical hypersensitivity. Together, these findings show that nerve injury functionally reallocates primary afferent drive away from inhibitory and toward excitatory spinal PVNs, establishing functional reallocation of afferent input as a mechanism of spinal disinhibition and a key determinant of mechanical allodynia.

## INTRODUCTION

The nervous system integrates sensory information into appropriate behaviours by distinguishing innocuous stimuli from aversive ones. In the somatosensory system, low-threshold mechanoreceptors (LTMRs) encode touch from the periphery ^1^. After nerve injury, the same Aβ low-threshold inputs can cause a painful response, a condition known as mechanical allodynia ^2,3^. This implies that perception is determined not only by the type of peripheral input, but by how that input is gated and integrated by central circuits.

The spinal dorsal horn is the first central site where primary afferent information is processed before ascending to supraspinal regions. Here, spinal inhibitory interneurons prevent tactile afferents from gaining access to the polysynaptic circuits that drive nociception ^4–7^. After injury, the loss of this inhibition opens the gate and permits mechanical allodynia ^8–13^. Mechanistic studies of this disinhibition have focused primarily on the inhibitory neurons themselves, their intrinsic excitability, and the reduction in their inhibitory output ^14–19^. However, these inhibitory interneurons can only gate sensory information if they are effectively recruited by sensory afferents in the first place. The nature of this recruitment and how it changes after nerve injury remain poorly understood. We therefore asked whether Aβ afferents continue to appropriately engage spinal inhibitory interneurons after nerve injury, and how the excitatory drive that Aβ fibers impose on these circuits is reorganized.

Parvalbumin-expressing interneurons (PVNs) are a well-defined element of this tactile gate, positioned between low-threshold afferents and polysynaptic nociceptive circuitry ^20–27^. Transient silencing of PVNs in naïve mice is sufficient to produce mechanical allodynia, whereas activating them relieves allodynia in nerve-injured mice ^20^. Because their behavioral role is established but the afferent drive they receive is not, PVNs offer a tractable entry point for asking how Aβ input is allocated onto spinal inhibitory circuits, and how nerve injury reorganizes this allocation.

Here, we ask how Aβ LTMR input is distributed across spinal PVNs and how this organization is altered after peripheral nerve injury. Using whole-cell patch-clamp electrophysiology in *ex vivo* spinal cord slices paired with dorsal root stimulation or optogenetics, together with pseudorabies circuit tracing and chemogenetic behavioural assays, we found that Aβ-LTMRs recruit PVNs through both direct excitation and feedforward inhibition. Unexpectedly, afferent stimulation evoked purely hyperpolarizing responses in a subset of PVNs, suggesting that some PVNs are themselves inhibited by Aβ input rather than simply relaying mixed excitatory/inhibitory drive. These results raise the possibility that the PVN population is not functionally homogeneous. To resolve this, we used intersectional genetic tools that have only recently become available to separate between inhibitory (iPVN) and excitatory subsets (ePVN) of PVNs. Under naïve conditions, Aβ input preferentially activates iPVNs, whereas ePVNs receive both direct excitation and Aβ-driven feedforward inhibition. After nerve injury, this drive is redistributed: iPVNs show reduced action potential recruitment, while ePVNs shift toward more excitatory responses. Silencing iPVNs in naïve mice produced nerve injury-like mechanical allodynia, whereas activating them after injury was analgesic. Conversely, activating ePVNs, whose behavioral role *in vivo* had not previously been tested, promoted hypersensitivity, and silencing them after nerve injury alleviated mechanical allodynia. We further identified a previously uncharacterized population of Complexin-1 (Cplx1)-expressing inhibitory interneurons as a candidate source of the Aβ-driven feedforward inhibition onto ePVNs. Together, these findings reveal that tactile gating depends on how Aβ-driven input is allocated across distinct, functionally opposed populations of spinal interneurons. We show that nerve injury can induce mechanical allodynia by shifting this allocation away from inhibitory and toward excitatory pathways.

## RESULTS

### The synaptic response profile of PVNs changes after nerve injury

We previously showed that dorsal horn PVNs are modality-specific to mechanical inputs from the periphery ^20^. Following nerve injury, a PVN-mediated loss of inhibitory tone in the dorsal horn has been consistently reported ^20,21,23–26^. Importantly, this disinhibition is due to functional deficits in PVNs rather than cell loss and has been attributed to decreased synaptic connectivity to their postsynaptic targets ^20^, weakened inhibitory output of PVNs ^23^, and decreased intrinsic excitability of PVNs ^21,24,25^. However, whether PVNs are appropriately recruited by peripheral primary afferents after nerve injury remains unknown. Without sufficient presynaptic activation of PVNs in the first place, downstream changes in their intrinsic excitability or post-synaptic efficacy become secondary. Moreover, these mechanisms are not mutually exclusive and may act in parallel to shape PVN function after nerve injury.

Here, we hypothesized that peripheral nerve injury reduces the recruitment of PVNs by Aβ-LTMRs, thereby decreasing their activation. We first confirmed the development of mechanical allodynia following chronic constriction injury (CCI) of the sciatic nerve by measuring mechanical nocifensive thresholds in naïve mice and at two post-injury time points: 5 days (CCI 5dp; acute phase) and 3 weeks (CCI 3wp; chronic phase). As expected, both CCI groups exhibited significantly reduced mechanical nocifensive thresholds compared with naïve mice (Figure 1Ai). Following behavioral assessment, we performed targeted whole-cell patch-clamp recordings from genetically labelled PVNs in transverse spinal cord slices of PVCre;tdTomato mice to examine Aβ-evoked synaptic responses (Figure 1Aii-iv). The dorsal root was kept intact and stimulated to activate primary afferents while recording PVN responses. In voltage clamp, all PVNs received monosynaptic Aβ input, and about 40% also received Aδ input, which remained unchanged in naïve and CCI 3wp (Figure S1A-C). Conduction velocity, latency of evoked excitatory post-synaptic current (eEPSC), and paired pulse ratio also were not statistically significant, although eEPSC amplitude was reduced at CCI 3wp (Figure S1D-H). These results show preserved primary afferent connectivity with reductions to synaptic strength after nerve injury. Next, we standardized the stimulation intensity across different slices by determining the stimulation intensity required to saturate Aβ-evoked responses based on previously reported protocols ^19,28^. The eEPSC amplitude did not continue to increase after 35 μA, so we used 75 μA and 4 Hz for all subsequent dorsal root stimulation experiments to ensure maximal Aβ fiber recruitment as this protocol did not engage other primary afferent inputs in our hands (Figure S2).

**Figure 1:**
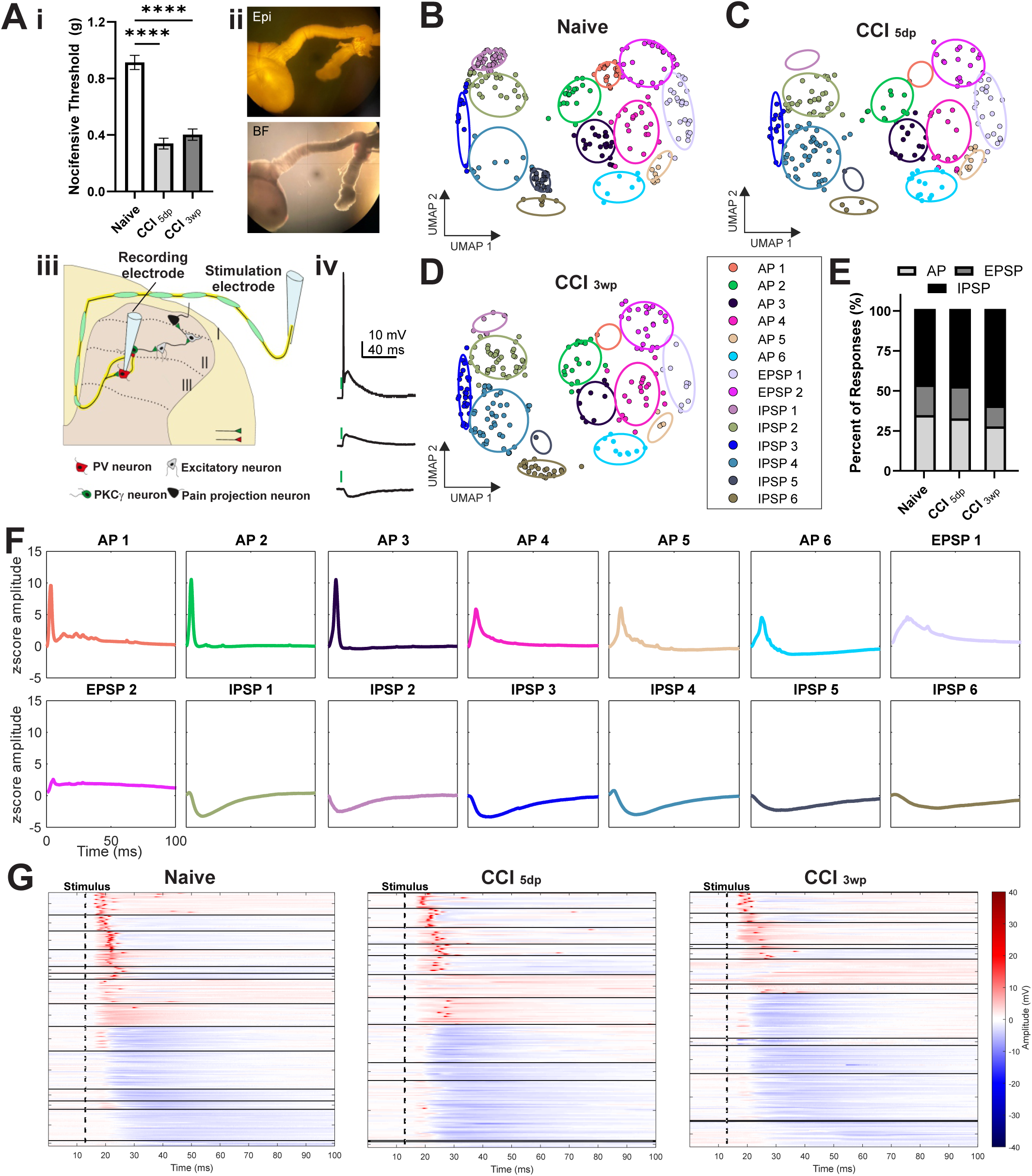
The synaptic response profile of PVNs changes after nerve injury A. (i) Mechanical nocifensive thresholds measured in naïve mice (n=23), at 5 days (5dp; n=14), and 3 weeks (3wp; n=17) following CCI surgery. One-way, ANOVA, Tukey’s multiple comparisons. (ii) Epifluorescence (Epi) and bright field (BF) image of the transverse spinal cord slice from PVcre;tdTomato mice, with the L5 dorsal root attached to a stimulating electrode. (iii) Schematic of the Aβ fiber-to-PVN circuit and recording configuration. (iv) Representative traces of the evoked responses of PVNs, action potential (AP, top), excitatory post-synaptic potential (EPSP, middle), inhibitory post-synaptic potential (IPSP, bottom). Stimulus is indicated by the green line on top of the traces. B-D. UMAP represents the individual synaptically evoked responses of PVNs from naïve (B; n=300 responses from 60 PVNs across 20 mice), 5dp-CCI (C; n= 200 responses from 40 PVNs across 12 mice), and 3wp-CCI (D; n=245 responses from 49 PVNs across 16 mice) conditions. Clusters are color-coded and delineated by ellipses. E. The percent of synaptically evoked responses of PVNs that responded with action potential (AP), excitatory post-synaptic potential (EPSP), and inhibitory post-synaptic potential (IPSP) in naïve, 5dp, 3wp mice. Chi-square contingency test, n.s. non-statistically significant. F. Mean synaptically evoked responses of each cluster and their classification. Amplitudes are shown as z-scored membrane potential (mV). G. Heatmap of each synaptically evoked response of PVNs organized by cluster for naïve, 5dp, 3wp conditions. Clusters are in chronological order as they appear in panel F.

We next assessed the functional output of PVNs to Aβ stimulation using current clamp. As the response types were heterogeneous, we used unbiased clustering analyses to visualize Aβ-evoked responses in PVNs using UMAP. Across conditions, evoked responses segregated into 14 clusters which belonged to action potential (AP), excitatory post-synaptic potentials (EPSP), and inhibitory post-synaptic potential (IPSP) response types (Figure 1B-D). In naïve mice, PVNs responses were split between excitatory (AP/EPSP) and inhibitory (IPSP) responses. This distribution was largely unchanged at CCI 5dp but shifted towards majority inhibitory responses at CCI 3wp (Figure 1E). Average cluster responses and population heatmaps showed the variation in response kinetics (Figure 1F-G). The population heatmaps revealed an increase in inhibitory response profiles at CCI 3wp (Figure 1G).

In the UMAPs, the responses did not cluster based off sex differences, batch effects across mice, or intrinsic firing (Figure S3A-C). Cluster-level analysis revealed selective remodeling of synaptic response types, including loss of some responses (e.g., AP1, IPSP5) after nerve injury, and expansion of others (e.g. IPSP6) during the chronic phase (Figure S3D). Cluster features when grouped by excitatory and inhibitory response types showed modest decrease in maximum response amplitude between naïve and CCI 3wp, and an increase in response integral between CCI 5dp and CCI 3wp (Figure S4). Notably, the number of Aβ synapses labelled by VGluT1/Homer1^29,30^ and miniature EPSCs (mEPSCs) parameters (Figure S5), as well as inhibitory synapses labelled with gephyrin^31^ and miniature IPSCs (mISPCs) parameters (Figure S6) remained unchanged after nerve injury. This data suggests that synapse loss or gain does not account for the changes we observed in Aβ-evoked PVN responses.

Together, these results reveal a noticeable reorganization of PVN synaptic response profiles following nerve injury, characterized by a shift toward increased inhibition despite preserved primary afferent input. Unexpectedly, Aβ fiber stimulation evoked robust inhibitory response types in almost half of the recorded PVNs. Therefore, we next sought to define the circuit mechanisms underlying these evoked inhibitory responses.

### PVNs receive Aβ-driven feedforward inhibition

To determine whether Aβ-evoked inhibitory responses in PVNs were mediated by synaptic inhibition, we pharmacologically blocked glycinergic and GABAergic transmission during evoked IPSPs (eIPSPs). Bath application of strychnine and/or bicuculline abolished eIPSPs, either by unmasking underlying excitatory responses or eliminating the inhibitory response entirely (Figure 2A–B and Figure S7A–B, respectively). This data confirms that these evoked responses are neurotransmitter-mediated inhibition.

**Figure 2:**
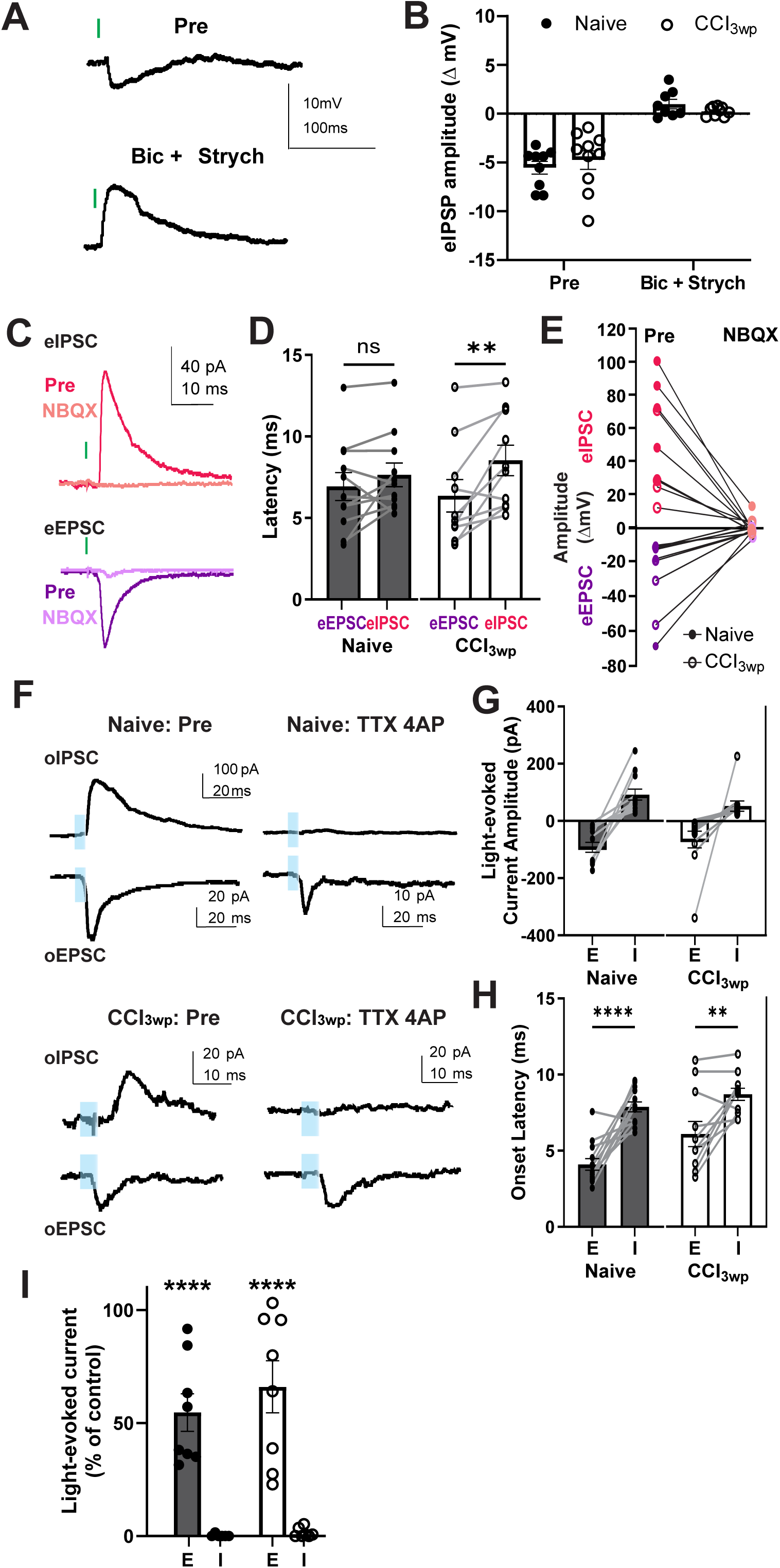
PVNs receive Aβ fiber-driven feedforward inhibition A-B. Representative current clamp traces (A) and quantification (B) of Aβ fiber synaptically evoked response onto PVNs before and after bicuculline (15μM) and strychnine (500nM) bath application in naïve (n=10 cells from 7 mice) and 3wp-CCI conditions (n=10 cells from 6 mice). C-E. Representative voltage-clamp traces (C), stimulus-to-response latency (D), and amplitude of evoked response (E) of Aβ fibers inputs onto PVNs. Electrically evoked IPSCs (eIPSCs; holding potential-40mV) and EPSCs (eEPSCs; holding potential-70mV) are shown before and after NBQX (10μM) bath application in naïve (n=5 cells from 2 mice) and 3wp-CCI (n=5 cells from 4 mice) conditions. Paired t-test. F-I. Representative voltage clamp traces from optogenetically evoked synaptic responses from primary afferents onto PVNs. Optically evoked IPSCs (oIPSCs;-40mV) and EPSCs (oEPSCs; - 70mV) are shown before and after TTX (1μM) and 4-AP (200μM) bath application in naive and 3wp-CCI mice (F). Quantification of light-evoked current amplitudes after TTX/4-AP application expressed as a percentage of baseline (I) are shown for naïve (n=8 cells from 2 mice) and 3wp-CCI (n=9 cells from 3 mice) conditions. Quantification of oEPSC and oIPSC amplitudes (G) and response onset latencies (H) are shown for naïve (n=12 cells from 3 mice) and 3wp-CCI (n=12 cells from 3 mice) conditions. Paired t-test.

We next investigated the circuit mechanism of this inhibition. We considered three possibilities: (1) co-release of GABA/glycine with glutamate from primary afferents, (2) axo-axonic synapse in which Aβ input recruit local inhibitory terminals that synapse onto PVNs, or (3) classical feedforward inhibition via an intermediate inhibitory interneuron. To test for co-release ^32^, we recorded eEPSCs and eIPSCs from the same PVN and blocked glutamatergic transmission with bath application of NBQX. This application abolished both the eEPSC and eIPSC in both naïve and CCI 3wp conditions (Figure 2C–E), indicating that the evoked inhibitory current depends on glutamate release and is not mediated by direct GABA/glycine release from primary afferents. We then distinguished between axo-axonic inhibition and polysynaptic feedforward inhibition. We expressed channelrhodopsin (ChR2) restricted to peripheral neurons in the dorsal root ganglion (DRG)^33^ through an intraperitoneal injection of AAV-php.S-CAG-ChR2-eGFP in neonatal mice (Figure S7C). This design allowed us to selectively photoactivate primary afferents terminals and record light-evoked responses in PVNs. To determine if the optically evoked IPSCs (oIPSCs) and EPSCs (oEPSCs) were the result of monosynaptic transmission, we blocked the propagation of action potentials through bath-application of tetrodotoxin (TTX) and 4-aminopyridine (4-AP).

This pharmacological manipulation revealed that oIPSCs were abolished, whereas oEPSCs persisted (Figure 2F–I and Figure S7D-F), indicating that the latter receive monosynaptic primary afferent inputs, whereas the former requires action potential propagation from an intermediate neuron, and is therefore polysynaptic. These results demonstrate that the eIPSPs elicited in PVNs by Aβ fibers are via polysynaptic feedforward inhibition. Consistent with electrical stimulation, photo-stimulation revealed a shift toward increased proportion of inhibitory response profiles after nerve injury, along with reduced response amplitude (Figure S7G–H). Notably, individual PVNs could exhibit different types of responses (AP, EPSP, IPSP) depending on the mode of Aβ fiber activation, either electrically or optically, indicating that recruitment of inhibitory pathways onto PVNs can depend on several factors, including the mode of activation, the coincident detection of inhibitory and excitatory inputs, among others (Figure S7I–K).

Together, these findings support a model in which Aβ afferents provide both direct excitatory input and indirect inhibition to PVNs via recruitment of an unknown population of inhibitory interneuron, forming a feedforward inhibitory circuit. Yet the question remains, why do only a subset of PVNs exhibit Aβ-evoked inhibition? Given the known heterogeneity of PVNs being both inhibitory and excitatory subtypes ^22,25^, we next asked whether PVN identity predicts response profile by Aβ activation.

### Reduced Aβ fiber activation of inhibitory PVNs after nerve injury

To specifically examine Aβ-evoked responses in inhibitory PVNs (iPVNs), we generated an intersectional GlyT2Cre;PVDre;Ai66D mouse strain to selectively label glycinergic iPVNs with tdTomato ^18,34,35^ (Figure 3A). Their spinal cord sections confirmed high specificity, with 86.9% of labeled cells expressing Pax2 and 84.5% expressing PV (Figure 3B).

**Figure 3:**
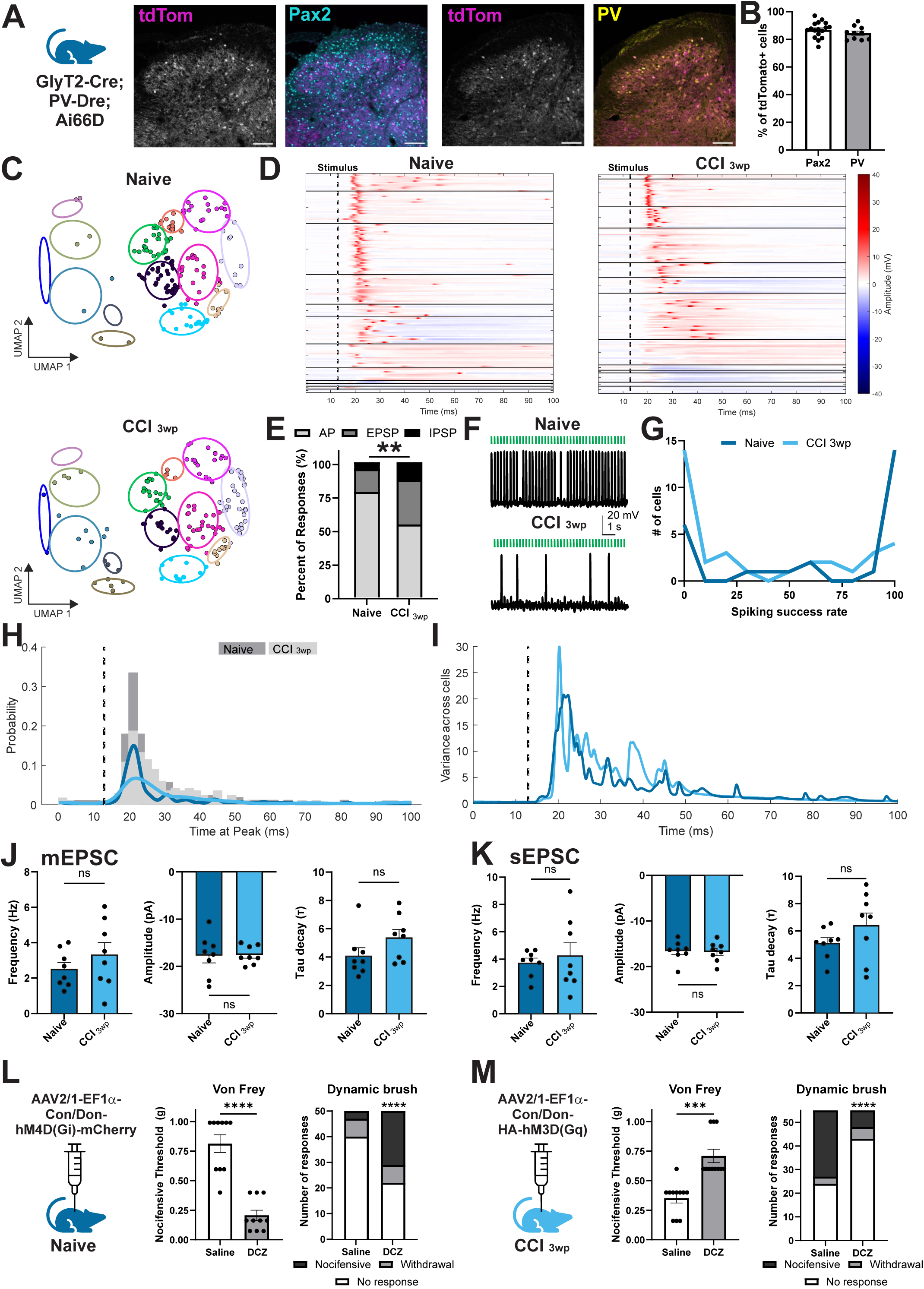
Reduced Aβ activation of inhibitory PVNs after nerve injury A-B. Inhibitory PVNs (iPVNs) are visually identifiable in a GlyT2Cre; PVDre; Ai66D mouse. Immunohistochemistry images of the dorsal horn and quantification (B) of tdTomato+ cells (magenta) that co-localize with Pax2 (cyan, white bar, 86.94 ± 1.39%, n=17 sections from 4 mice) and PV (yellow, grey bar, 84.51 ± 1.52%, n=10 sections from 3 mice). Scale bar 100μm. C. UMAP represents the individual synaptically evoked responses of iPVNs from naïve (top; n=155 responses from 31 iPVNs across 8 mice), and 3wp-CCI (bottom; n=165 responses from 33 PVNs across 7 mice) conditions. Clusters are color-coded and delineated by ellipses. D. Heatmap of each synaptically evoked response of iPVNs organized by cluster for naïve, 3wp conditions. E. The percent of synaptically evoked responses of iPVNs that responded with action potential (AP), excitatory post-synaptic potential (EPSP), and inhibitory post-synaptic potential (IPSP) in naïve, 3wp mice. Chi-square contingency test, **p-value=0.0015. F-G. Representative traces (F) of iPVNs in naïve (top) and 3wp-CCI (bottom) mice in response to 100 Aβ-fiber stimulations (4Hz, 75μA, 25 seconds). Panel G shows the quantification of the spiking success rate out of 100 stimulations in naïve (dark blue, n=26 cells from 8 mice) and 3wp-CCI (light blue, n=33 cells from 7 mice). H. Time at peak probability distribution of iPVNs in naïve (dark grey bars) and 3wp-CCI (light grey bars) fitted with Kernel density estimation. I. Variance across Aβ-evoked responses in iPVNs across time in naïve and 3wp-CCI. J. The frequency, amplitude, tau decay of mEPSCs in iPVNs of naïve (n=8 cells from 3 mice) and 3wp-CCI (n=9 cells from 4 mice). Unpaired t-test. K. The frequency, amplitude, tau decay of sEPSCs in iPVNs of naïve (n=8 cells from 3 mice) and 3wp-CCI (n=8 cells from 4 mice). Unpaired t-test. L. Naïve GlyT2Cre;PVDre;Ai66D mice were injected with AAV2/1-EF1α-Con/Don-hM4D(Gi)-mCherry. Static mechanical nocifensive threshold (von Frey, left, paired t-test, ****p-value<0.0001) and distribution of behavioural responses to dynamic mechanical brush stimulations (right, Chi-square contingency test, ****p-value<0.0001) were measured after saline (n=10 mice) or DCZ (n=10 mice) treatment. M. GlyT2Cre;PVDre;Ai66D mice were injected with AAV2/1-EF1α-Con/Don-HA-hM3D(Gq). At 3 weeks post-CCI, static mechanical nocifensive threshold (von Frey, left, paired t-test, ***p-value=0.0002) and distribution of behavioural responses to dynamic mechanical brush stimulations (right, Chi-square contingency test,****p-value<0.0001) were measured after saline (n=11 mice) or DCZ (n=11 mice) treatment.

Next, we recorded from iPVNs during Aβ fiber stimulation in naïve and at the chronic time point CCI 3wp, as no significant changes to response distributions were observed at CCI 5dp (Figure 1E). The evoked responses were projected into the same UMAP space defined in Figure 1 (Figure 3C). Notably, iPVNs predominantly responded with excitatory responses (AP/EPSP) and rarely exhibited inhibitory response profiles when activated by primary afferent inputs, consistent with their identity as inhibitory output neurons (Figure 3D). Following nerve injury, iPVNs displayed a change in their evoked responses. There was a reduction in action potential firing and a relative increase in subthreshold EPSPs, indicating weakened functional recruitment of iPVNs by Aβ fibers. Overall, this balance shifted toward reduced excitatory drive and a reduction in the proportion of iPVNs responding with APs (Figure 3E). To directly assess primary afferent recruitment fidelity, we applied a train of 100 Aβ stimulations (4 Hz; 25 seconds). In naïve mice, iPVNs reliably generated action potentials with high success rates, whereas in CCI 3wp mice, failure rates increased significantly (Figure 3F–G, Figure S8). Moreover, spike timing became significantly more variable after nerve injury, with broader and flatter distributions of peak response times (Figure 3H) and increased across-cell variability (Figure 3I), indicating degraded temporal precision of Aβ-driven activation of iPVNs. Next, we measured mEPSCs and sEPSCs to examine if the contribution from Aβ inputs were enough to produce a significant change in global excitatory input after nerve injury. Notably, we found that mEPSC and sEPSC frequency, amplitude, and decay properties remained unchanged (Figure 3J–K).

To test the causal contribution of iPVNs to sensory gating, we bidirectionally manipulated their activity *in vivo* using intersectional chemogenetics. Selective silencing of iPVNs using a Cre/Dre double-dependent AAV encoding the inhibitory chemogenetic receptor, hM4D-Gi ^36^, reduced nocifensive mechanical thresholds and increased nocifensive responses to dynamic brush, indicating mechanical hypersensitivity (Figure 3L), similar to previous ablation experiments ^35^. Conversely, activation of iPVNs using the excitatory chemogenetic receptor, hM3D-Gq, in CCI 3wp mice alleviated mechanical allodynia, increasing nocifensive mechanical threshold to von Frey filaments and reducing dynamic brush-evoked nocifensive responses (Figure 3M).

Together, these results demonstrate that iPVNs are critical gatekeepers of touch and that their recruitment by Aβ afferents predominantly elicits action potentials. However, after nerve injury, these evoked excitatory responses are impaired, leading to the development of mechanical allodynia. This reduction in evoked excitatory responses occurs despite no significant changes to global synaptic input (Figure 3J-K), suggesting an increase in failure rate of circuit recruitment after nerve injury.

### Excitatory PVNs receive feedforward inhibition from Aβ fiber activation

We showed previously that a large portion of PVNs also respond with eIPSPs when recruited by Aβ fibers. Thus, we postulate that these eIPSPs could be occurring in excitatory PVNs (ePVNs). To specifically examine Aβ-evoked responses in ePVNs, we generated an intersectional VGluT2Cre;PVDre;Ai66D mouse strain to selectively label glutamatergic ePVNs with tdTomato ^34,37^ (Figure 4A). We validated the specificity of this strategy by immunohistochemistry approaches and observed that 83.1% of tdTomato-labeled neurons were Tlx3-positive and 85.3% were PV-positive (Figure 4B).

**Figure 4:**
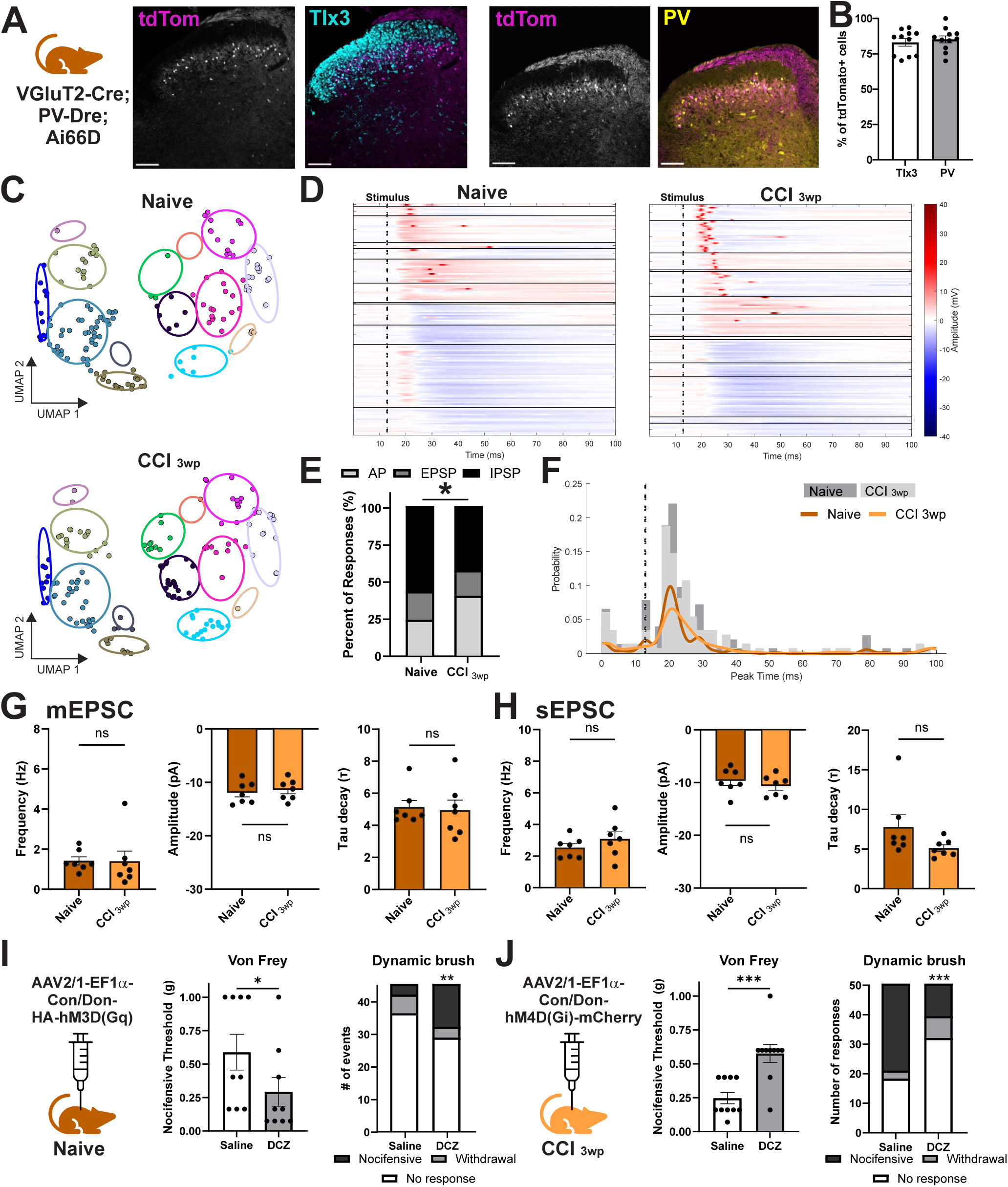
Excitatory PVNs receive feedforward inhibition from Aβ fiber activation A-B. Excitatory PVNs (ePVNs) are visually identifiable in a VGluT2Cre; PVDre; Ai66D mouse. Immunohistochemistry images of the dorsal horn and quantification (B) of tdTomato+ cells (magenta) that co-localize with Tlx3 (cyan, white bar, 83.11± 2.65%, n=11 sections from 3 mice) and PV (yellow, grey bar, 85.3 ± 2.48%, n=11 sections from 3 mice). Scale bar 100μm. C. UMAP represents the individual synaptically evoked responses of ePVNs from naïve (top; n=155 responses from 31 ePVNs across 8 mice), and 3wp-CCI (bottom; n=150 responses from 30 PVNs across 6 mice) conditions. Clusters are color-coded and delineated by ellipses. D. Heatmap of each synaptically evoked response of ePVNs organized by cluster for naïve, 3wp conditions. E. The percent of synaptically evoked responses of ePVNs that responded with action potential (AP), excitatory post-synaptic potential (EPSP), and inhibitory post-synaptic potential (IPSP) in naïve, 3wp mice. Chi-square contingency test, *p-value=0.048. F. Time at peak probability distribution of ePVNs in naïve (dark grey bars) and 3wp-CCI (light grey bars) fitted with Kernel density estimation. G. The frequency, amplitude, tau decay of mEPSCs in ePVNs of naïve (n=7 cells from 3 mice) and 3wp-CCI (n=7 cells from 3 mice). Unpaired t-test. H. The frequency, amplitude, tau decay of sEPSCs in ePVNs of naïve (n=7 cells from 3 mice) and 3wp-CCI (n=7 cells from 3 mice). Unpaired t-test. I. Naïve VGluT2Cre;PVDre;Ai66D mice were injected with AAV2/1-EF1α-Con/Don-HA-hM3D(Gq). Static mechanical nocifensive threshold (von Frey, left, paired t-test, *p-value=0.0124) and distribution of behavioural responses to dynamic mechanical brush stimulations (right, Chi-square contingency test,**p-value=0.0091) were measured after saline (n=9 mice) or DCZ (n=9 mice) treatment. J. VGluT2Cre;PVDre;Ai66D mice were injected with AAV2/1-EF1α-Con/Don-hM4D(Gi)-mCherry. At 3 weeks post-CCI, static mechanical nocifensive threshold (von Frey, left, paired t-test, ***p-value=0.0004) and distribution of behavioural responses to dynamic mechanical brush stimulations (right, Chi-square contingency test,***p-value<0.0004) were measured after saline (n=10 mice) or DCZ (n=10 mice) treatment.

Evoked responses from ePVNs during Aβ stimulation revealed a prominently more diverse response profile than from iPVNs. When mapped into the same UMAP space, ePVNs distributed across both excitatory and inhibitory clusters (Figure 4C–D), with approximately half of responses consisting of eIPSPs. This distribution indicates that Aβ recruitment of ePVNs can engage strong secondary inhibitory inputs. Unlike iPVNs, the overall proportion of response types in ePVNs was shifted towards increased excitatory responses after nerve injury (Figure 4E), including increased action potential prominence and reduced inhibitory responses (Figure 4D). Similar to iPVNs, ePVNs exhibited degraded temporal precision after injury, with broader response timing distributions (Figure 4F). Baseline excitatory synaptic properties also remained unchanged (Figure 4G–H), as in iPVNs. Despite their known presence in the dorsal horn ^14,22,25^, the contribution of these ePVNs have never been examined *in vivo*. We set out to test their functional role with chemogenetic manipulation. Activation of ePVNs (hM3Dq) increased mechanical sensitivity, lowering nocifensive thresholds and enhancing nocifensive responses to brush dynamic (Figure 4I). In contrast, silencing ePVNs alleviated mechanical allodynia in CCI 3wp mice (Figure 4J), assessed using von Frey filaments and dynamic brush assays.

Together, these results identify ePVNs as a previously unappreciated neuronal population of dorsal horn touch circuits. They receive Aβ excitatory input, like their inhibitory counterparts, but this excitatory input is masked by a secondary recruitment of inhibition onto these neurons, not only preventing them from firing, but also eliciting eIPSPs. Unlike iPVNs, their recruitment by Aβ afferents is preserved after injury, suggesting that dorsal horn circuitry imbalance arises from reduced iPVN activation, and hence inhibitory output, combined with maintained or functionally redirected excitatory output from ePVN activity.

### Divergent intrinsic and circuit properties distinguish iPVNs and ePVNs

To further compare these two populations of PVNs, we examined intrinsic firing and synaptic properties. iPVNs displayed adapting firing patterns, whereas ePVNs exhibited sustained, tonic firing with higher spike frequencies (Figure S8A–B, Supplementary Table 1), contrary to classical expectations ^38–40^. Despite these differences, both populations showed reduced spike output after nerve injury, which we had shown previously ^25^. Aβ-driven synaptic properties, like paired-pulse ratio and eEPSC latency were unchanged across groups (Figure S8C–E). Evoked current amplitude was significantly lower in ePVN Naïve compared to iPVN Naive and in iPVN CCI 3wp compared to iPVN Naive (Figure S8F), consistent with our current clamp recordings. No sex or batch effects were observed (Figure S9A–C). However, cluster-level analyses revealed strong cell-type-specific biases. For example, AP1 responses were enriched in iPVNs, whereas IPSP clusters were preferentially elicited in ePVNs (Figure S9D). Within AP response types, ePVNs show reduced peak amplitudes relative to iPVNs, and in CCI 3wp exhibited increased minimum amplitudes compared to both naïve ePVNs and iPVNs (Figure S10). mEPSC and sEPSC parameters, half-width, rise time, decay time, and coefficient of variation of the peak amplitude, showed no differences between iPVNs and ePVNs in naïve and CCI 3wp mice (Figure S11). Behaviorally, chemogenetic manipulation of either population did not alter thermal sensitivity in Hargreaves’ or acetone assays, indicating modality specificity to mechanical, but not thermal, processing ^20^ (Figure S12).

Together, these data demonstrated a functional difference between iPVNs and ePVNs. We showed that iPVNs lose effective Aβ-driven recruitment after nerve injury, whereas ePVNs remain recruited, have a higher proportion of AP responses, and a lower proportion of IPSPs. Furthermore, the identity of the neurons responsible for the Aβ-fiber driven feedforward inhibition onto ePVNs remains to be determined. The source of their inhibitory input is likely to be a key determinant of circuit function, shaping the timing, routing, and downstream impact of Aβ-evoked activity.

### A subset of Complexin-1-expressing interneurons mediates Aβ-driven feedforward inhibition onto ePVNs

PVNs are known to inhibit other PVNs across multiple CNS regions ^41,42^, including the spinal dorsal horn ^22^. Therefore, we first asked whether the Aβ-driven eIPSPs observed in PVNs is due to PVN–PVN inhibition. To examine this possibility, we used PVCre;ChR2-tdTomato mice ^43^ to selectively photoactivate PVNs (Figure S13A) and performed two complementary electrophysiological protocols. First, we assessed whether Aβ stimulation recruits PVN-mediated inhibition onto PVNs by combining electrical dorsal root Aβ fiber stimulation with photostimulation of PVNs. If Aβ activation engages PVN–PVN inhibition, co-stimulation with blue light should occlude the evoked inhibitory current. However, simultaneous electric-and photo-stimulation did not alter Aβ-evoked IPSC amplitude in either naïve or CCI 3wp conditions (Figure 5A–B), arguing that either PVN-PVN inhibition is already recruited by Aβ stimulation, or that PVN-PVN inhibition is not present in that circuit. Next, we tested the presence of Aβ-driven PVN-PVN inhibition more directly using synaptic depletion. Prolonged photostimulation was used to deplete PVN-mediated inhibitory transmission, confirmed by a reduced response to a subsequent light pulse (Figure 5C top panels, Figure 5D). If Aβ-evoked IPSPs were mediated by PVNs, this depletion should reduce or abolish the electrically-stimulated Aβ-evoked inhibitory current. Instead, electrically-stimulated Aβ-evoked IPSCs remained intact following photo-depletion (Figure 5C middle panels, Figure 5D, Figure S13B), demonstrating that the feedforward inhibition is not mediated by PVNs but rather by a separate inhibitory interneuron population.

**Figure 5:**
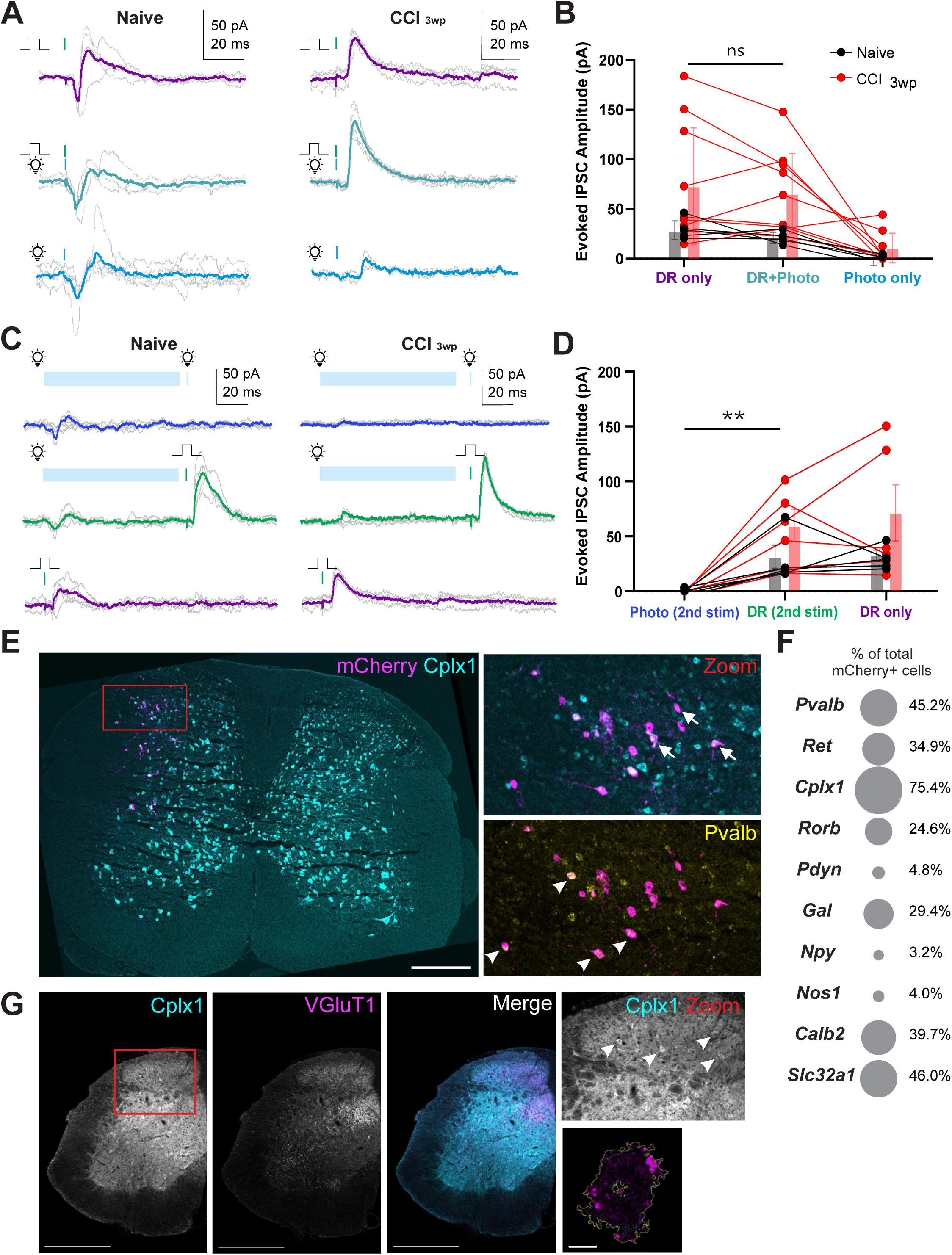
A subset of Complexin-1-expressing interneurons mediates Aβ-driven feedforward inhibition onto ePVNs A-B. Representative voltage-clamp traces (A) and quantification (B) of a PVN from a PVcre;ChR2-tdTomato mouse in response to Aβ fiber stimulation (top), Aβ fiber and photo-stimulation (middle), and photo-stimulation alone (bottom). PVNs were held at-40mV. Two-way ANOVA, Šídák’s multiple comparison. C-D. Representative voltage-clamp traces (A) and quantification (B) of a PVN from a PVcre;ChR2-tdTomato mouse in response to a 90-ms depletion photo-stimulation followed by a second photo pulse (top), a 90-ms depletion photo-stimulation followed by an Aβ fiber electrical stimulation (middle), a single Aβ fiber electrical stimulation (bottom). PVNs were held at - 40mV. Two-way ANOVA, Šídák’s multiple comparison,**p-value=0.0034. E. Representative image of fluorescent *in situ* hybridization of rabies labelled pre-synaptic neurons (mCherry+ magenta cells) to PVNs (yellow, white arrowheads) colocalize with pre-synaptic Complexin-1 (Cplx1)-expressing cells (cyan, white arrows). Scale bar 500μm. F. Quantification of the percentage of the colocalization of different neuron markers with the pre-synaptic mCherry+ neurons of PVNs. n=125 mCherry+ cells from 2 mice. G. Representative image of immunohistochemistry of VGluT1 (magenta)-positive punta colocalized on Cplx1 (cyan)-expressing cells. Scale bar 500μm, 80mm. Bottom right image shows a reconstructed Cplx-positive cell outlined in yellow and the colocalized VGluT1-positive punta. 6.3±0.74 VGluT1-positive punta per Cplx1-positive cell, n=27 cells from 3 mice.

To identify these presynaptic inhibitory neurons, we performed retrograde trans-synaptic rabies tracing from PVNs. Consistent with our electrophysiological findings (Figure S1), fluorescent *in situ* hybridization on DRG tissue confirmed monosynaptic input from myelinated Aβ afferents (Nefh+/Graf1+) ^44,45^ in both naïve and CCI 3wp conditions (Figure S13C). Within the spinal cord, a substantial portion of presynaptic neurons was inhibitory (Pax2+; 33% naïve, 36% CCI 3wp), indicating a prominent inhibitory connection upstream of PVNs (Figure S13D-F). As the inhibitory responses were predominantly found in ePVNs, we performed retrograde trans-synaptic rabies tracing in the VGluT2-Cre;PV-Dre mouse. Then, to molecularly define this presynaptic inhibitory population, we performed fluorescent *in situ* hybridization targeting canonical inhibitory neuron markers ^46–48^. Among these, Complexin-1 (Cplx1) showed a higher proportion of colocalization with rabies-labeled presynaptic neurons (Figure 5E-F; Figure S14), identifying these inhibitory neurons as candidate mediators of this circuit. Consistent with a role in Aβ-driven inhibition of ePVNs, Cplx1+ neurons received VGluT1+ appositions (6.3 ± 0.7 VGluT1-positive puncta per Cplx1-positive soma), indicative of direct input from myelinated low-threshold mechanoreceptors, like Aβ fibers ^29^ (Figure 5G). Together, these findings identify a previously uncharacterized population of Cplx1-expressing inhibitory interneurons that likely mediate Aβ-driven feedforward inhibition onto ePVNs. These results identify a new circuit for how tactile input engages inhibitory control of excitatory interneurons within the dorsal horn to gate touch-evoked pain.

## DISCUSSION

Spinal inhibition of touch-evoked pain proposes that inhibitory interneurons prevent innocuous inputs from engaging central nociceptive circuits and that mechanical allodynia emerges when this inhibition is reduced after nerve injury. Understanding the mechanism for how this disinhibition occurs can reveal novel therapeutic insights to neuropathic pain. Previous work has primarily focused on changes in inhibitory neuron intrinsic firing and their downstream synaptic impact on excitatory interneurons after nerve injury. In this study, we studied the recruitment of interneurons by primary afferents and identified a previously unrecognized level of regulation in the dorsal horn that contributes to disinhibition and mechanical allodynia. We showed that Aβ-LTMR input is not distributed uniformly across dorsal horn PVNs, but instead engages them differentially according to cell identity, defined as inhibitory (iPVNs) and excitatory PVNs (ePVNs). Following nerve injury, the Aβ-driven input onto PVNs becomes bidirectionally reorganized. Aβ afferents become less efficient at recruiting iPVNs yet more efficient at recruiting ePVNs. Together, these findings suggest that the disinhibition underlying mechanical allodynia arises not only from a reduction in inhibition, but from a reversal of peripheral afferent input engagement that simultaneously weakens inhibitory interneuron recruitment while strengthening excitatory ones.

We report significant changes in spinal circuit recruitment after CCI despite relatively preserved anatomical connectivity. A notable finding was that nerve injury caused the overall PVN population to shift toward inhibitory responses to Aβ stimulation (Figure 1E), which appears inconsistent with the decrease in IPSP in ePVNs (Figure 4E), despite an increase in IPSP in iPVNs (Figure 3E). However, these observations are due to approximately two-thirds of PVNs being inhibitory, whereas one-third are excitatory ^25^, thus recordings from the pan-PVN population are disproportionately influenced by changes occurring within the iPVNs. Further, monosynaptic input to PVNs remained intact after CCI, and measures of afferent connectivity, including conduction velocity, response latency, paired-pulse ratio, and synapse density were largely unchanged (Figure S1, S5, and S6). Spontaneous excitatory synaptic properties also were unchanged in both iPVNs and ePVNs (Figure 3K, 4H, and S11). Further, since sEPSCs are shaped by multiple factors including presynaptic firing rate, release probability, and ongoing network activity^49^, the absence of change in sEPSCs can also reflect compensatory changes that offset a genuine Aβ-specific deficit. Nevertheless, Aβ stimulation evoked markedly different response profiles in PVNs after nerve injury. These observations demonstrate that structural connectivity alone is insufficient to predict circuit function and suggest that nerve injury induces changes in how existing circuits are engaged rather than rewiring of sensory pathways. This reallocation of sensory recruitment of spinal interneurons may help reconcile previous reports of modest anatomical changes with the drastic behavioural deficits observed in neuropathic pain ^50–52^. Our data supports the conclusion that large-scale anatomical rewiring is unlikely, but does not exclude finer structural plasticity, such as changes in synapse size, receptor composition, or subcellular/dendritic location, as a contributor to the functional changes reported. Our findings also refine our previous characterization of PVN intrinsic excitability after nerve injury. In our previous study examining pan-PVNs^25^, tonic firing was attributed to inhibitory PVNs based on prevailing assumptions regarding dorsal horn neuron firing^38–40,53^. By genetically resolving inhibitory and excitatory PVN populations, we demonstrated that tonic firing is instead a feature of ePVNs, whereas iPVNs predominantly exhibited initial burst followed by rhythmic firing (Figure S8A-B). Importantly, this revised classification does not alter the principal conclusion of our previous work, as both populations exhibit reduced intrinsic excitability and frequency adaptation following nerve injury.

Our results further refine current understanding of iPVNs as tactile gatekeepers of the dorsal horn. Consistent with previous studies, we found that silencing iPVNs induced mechanical allodynia ^20,35^ and that they were preferentially activated by Aβ input under naïve conditions. We found that nerve injury selectively reduced the reliability of Aβ-driven action potential generation and greater temporal variability despite largely preserved baseline synaptic properties (Figure 3). mEPSCs and sEPSCs reflect the aggregate activity of all excitatory inputs converging onto a neuron and do not distinguish primary afferent synapses from other local or long-range excitatory sources. Consequently, changes restricted to the Aβ-iPVN synapse may be diluted and therefore remain undetectable in global measures of spontaneous synaptic transmission. Notably, we do observe decreased Aβ-evoked EPSC amplitude on iPVNs at CCI 3wp (Figure S8F). Together, these findings suggest that tactile gating depends not only on the inhibitory output of iPVNs, but also on their effective recruitment by sensory afferents. In this framework, iPVNs can remain structurally connected yet become functionally disengaged from incoming sensory inputs. Such a failure of recruitment represents a previously underappreciated mechanism contributing to dorsal horn disinhibition.

We also identified an Aβ-driven feedforward inhibitory circuit onto around 50% of PVNs, revealing that primary afferents simultaneously engage both excitation and secondary inhibition within dorsal horn networks. Although the *in vitro* pharmacological and optogenetic experiments defining this feedforward circuit were performed in PVCre;tdTomato mice, these analyses were restricted to PVNs that exhibited Aβ-evoked IPSP responses (Figure 2). Subsequent cell-type-specific recordings revealed that such inhibitory response profiles were rare among iPVNs yet prominent in ePVNs, where more than half of recorded excitatory neurons exhibited eIPSPs following Aβ stimulation (Figure 3D-E and Figure 4D-E). Thus, while the initial experiments were performed in a pan-PVN population, their response characteristics strongly suggest that the majority of neurons receiving the feedforward inhibitory response were ePVNs. Pharmacological and optogenetic manipulations demonstrated that these inhibitory responses arise through a polysynaptic glutamate-dependent mechanism rather than neurotransmitter co-release from primary afferents or direct axo-axonic inhibitory transmission (Figure 2). An unexpected feature of the feedforward inhibition we observed was its precise temporal organization. In the naïve condition, the onset latency of the eIPSC was not significantly longer than the direct eEPSC (Figure 1A-iv and Figure 2D), despite the eIPSC requiring an additional synapse. This observation suggests that the circuit is organized to compensate for the additional synaptic delay inherent in feedforward inhibition. Although the underlying mechanism remains unknown, several possibilities could account for this compensation. For example, Aβ afferents contacting the feedforward inhibitory neuron may exhibit faster conduction velocity or shorter path lengths than those providing direct excitation ^1,54^. Alternatively, the secondary inhibitory neuron themselves may possess intrinsic excitable properties optimized for rapid action potential generation or propagation and minimize synaptic delay ^55,56^. Notably, after nerve injury, this near-synchronous arrival of eEPSC and eIPSC is impaired, with the eEPSC arriving significantly earlier (Figure 2D). This loss of temporal precision and the decrease in eEPSC amplitude (Figure S1F) provides a possible explanation for the shift toward evoked excitatory responses observed in ePVNs after nerve injury. Thus, mechanical allodynia may arise from the disruption of the temporal coordination of excitatory and inhibitory pathways.

This circuit organization resembles canonical feedforward inhibitory motifs found throughout the central nervous system, in which sensory excitation recruits inhibitory interneurons that inhibit downstream network activity ^57^. Such circuits regulate gain, sharpen temporal fidelity, and restrict abnormal excitation ^41,55^. The presence of primary afferent driven feedforward inhibition on both inhibitory and excitatory interneurons has been previously described in the dorsal horn and has been proposed to balance the excitability of neurons, in the case of inhibitory neurons, and gate touch inputs from activating pain circuits, in the case of excitatory neuron ^19,58^. Our findings support this role but further suggest a distinct functional interpretation. Rather than limiting neural excitation, we propose that feedforward inhibition may act as a circuit-level routing mechanism that determines how sensory drive is distributed among parallel dorsal horn pathways. The differential organization of Aβ-drive onto PVNs places feedforward inhibition in a strategic position to regulate the relative engagement of iPVNs and ePVNs. Under naïve conditions, such inhibition may bias tactile afferent drive toward iPVNs while limiting activation of ePVNs, thereby preventing the activation of nociceptive circuits. Following nerve injury, the shift toward increased recruitment of ePVNs suggests that this balance is altered, allowing tactile input to be redistributed toward excitatory pathways. Our findings suggest that feedforward inhibition may selectively regulate the allocation of Aβ sensory routing within spinal circuits.

Importantly, this feedforward inhibition was almost entirely concentrated within the ePVN population, making up about 50% of their responses to Aβ stimulation (Figure 4E). These results suggest ePVNs act not only as excitatory relays of touch information, but as neurons whose activation is under tight inhibitory control. Behavioural experiments support a pro-allodynic role for ePVNs, as chemogenetic activation caused mechanical hypersensitivity, whereas silencing alleviated mechanical allodynia after nerve injury (Figure 4I-J). Together, these findings suggest a model where Aβ inputs engage at least two competing parallel pathways within the dorsal horn: (1) one that recruits iPVNs to prevent touch inputs from activating pain circuits, and (2) another that recruits ePVNs but simultaneously tunes their activity through feedforward inhibition. Following CCI, reduced activation of iPVNs combined with increased excitation of ePVNs may push the systems toward increased global excitation, thereby facilitating erroneous access of touch inputs to nociceptive pathways. The downstream consequence of increased ePVN excitation after CCI is also an important consideration. Previous studies showed that PVNs do not receive excitatory input from other ePVNs ^22^. Using optogenetic activation of PVNs in the presence of glycine and GABA receptor antagonists to isolate the ePVN population, Gradwell and colleagues demonstrated that photostimulation could only evoke action potentials after inhibition was unmasked. Consistent with this observation, they further reported that 12-36% of lamina I projection neurons received monosynaptic excitatory input from ePVNs, while 22% received polysynaptic input ^22^. Together with our behavioural findings, these results support a model in which ePVNs form part of an excitatory pathway capable of engaging nociceptive transmission and that their activity is under tight inhibitory control. The shift toward increased excitatory recruitment of ePVNs following CCI would therefore grant innocuous Aβ inputs access to these nociceptive pathways.

We also identified a subset of inhibitory Cplx1-expressing neurons as candidate mediators of Aβ-driven feedforward inhibition on ePVNs. Although dorsal horn inhibitory populations have been extensively catalogued by transcriptomics ^47,48^, the circuit functions of many molecularly defined subtypes remain poorly understood. Our rabies tracing and anatomical data indicate that inhibitory Cplx1-expressing neurons receive direct LTMR input and form monosynaptic synapses to ePVNs (Figure 5E-G), identifying them as strong candidate components rather than definitive mediators of this feedforward circuit. This population has not been previously implicated in touch-evoked pain circuits. We postulate that these Cplx1-expressing neurons may regulate the timing and distribution of Aβ-evoked activity within dorsal horn circuits. Whether these Cplx1-expressing neurons preferentially target ePVNs, participate in boarder inhibitory networks, or undergo nerve injury-induced plasticity remains to be further investigated.

In summary, our findings reveal a novel organization of dorsal horn circuitry in which Aβ afferent inputs are distributed through parallel pathways that differentially engage iPVNs and ePVNs. Peripheral nerve injury selectively disrupts recruitment of iPVNs, while increasing activation of ePVN-associated pathways, producing an imbalance that favours pathological activation of nociceptive circuits by touch inputs. Our results supplement the classic disinhibition hypothesis ^59–61^ by adding a previously unrecognized upstream recruitment mechanism that may improve our understanding of changes in dorsal horn circuitry after nerve injury. More broadly, our findings highlight a principle of circuit organization known as degeneracy, in which a neural system can contain parallel pathways that receive similar inputs yet generate distinct behavioral outputs ^62,63^. We suggest that feedforward inhibition can serve not only as a mechanism for controlling excitability but also as a mechanism for allocating information between competing circuit states. Thus, pathological states may not arise from the formation of new circuits or the loss of existing ones but perhaps can emerge from altered recruitment of pre-existing pathways. Therefore, circuit reallocation like the one we propose in this paper may represent a general principle by which neural networks achieve both flexibility and robustness in sensory processing, learning, and behavior. Understanding how information is dynamically routed through parallel neural circuits may therefore provide a useful framework for studying state-dependent computation across the nervous system.

## MATERIALS AND METHODS

### Mice

Mice were kept on a 12-h:12-h light/dark cycle, with food and water provided ad libitum. Both male and female mice were used for all experiments unless otherwise indicated. All experimental procedures were approved by the Animal Care and Use Committee at McGill University, in accordance with the regulations of the Canadian Council on Animal Care. PVCre, Ai14 (Cre dependent tdTomato reporter), GlyT2Cre, VGluT2Cre, Ai27D (Cre dependent ChR2) were obtained from Jackson Laboratory (#008069, #017320, #038515, #016963, #012567, respectively). PVDre and Ai66D (Dre and Cre dependent tdTomato reporter) were generated by the lab of Dr. Zeilhofer. PVCre, PVCre;tdTomato, PVDre;GlyT2Cre;Ai66D, PVDre;VGluT2Cre;Ai66D, PVDre;GlyT2Cre, PVDre;VGluT2Cre, PVCre;ChR2 hybrids were generated and maintained by in-house breeding.

### Fluorescent *in situ* hybridization

Mice were deeply anesthetized with 5% isoflurane and euthanized by cervical dislocation. DRGs were rapidly dissected and embedded in Optimal Cutting Temperature compound (OCT) and flash frozen on dry ice. Transverse sections (14 µm) were cut using the cryostat (Lecia Microsystems), directly mounted on Fisherbrand Superfrost Plus Microscope Slides (Cat. No. 12-550-15) and stored on dry ice. RNAscope fluorescent in situ hybridization was performed using the RNAscope® Multiplex Fluorescent Reagent Kit V2 (Advanced Cell Diagnostics, #323100), a TSA-based Fluorescent RNAscope®detection assay with 4-plex capability, in combination with the RNAscope 4-Plex Ancillary Kit for Multiplex Fluorescent v2 (#323120), or the RNAscope® Intro Pack for HiPlex12 Reagents Kit (488, 550, 650) v2 (#324443). All procedures were performed according to the manufacturer’s Multiplex Fluorescent v2 user manual or the HiPlex Assay v2 user manual. Tissue sections were treated with Protease IV (#322381) and hybridized with the following RNAscope probes: mCherry (#431201), Mm-Nefh-C2 (#443671), and Mm-Graf1-C4 (#31781), and the following HiPlex probes: mCherry-T1 (#431201), Mm-Pvalb-T2 (#421931), Mm-Ret-T4 (#431791), Mm-Cplx1-T5 (#482531), Mm-Rorb-T6 (#444271), Mm-Pdyn-T7 (#318771), Mm-Gal-T8 (#400691), Mm-Npy-T9 (#313321), Mm-Nos1-T10 (#437651), Mm-Calb2-T11 (#313641), Mm-Slc32a1-T12 (#319191). Control slides were run simultaneously and treated with a mouse RNAscope® 4-Plex Positive Control Probe-Mm (#321811) and RNAscope® 4-Plex Negative Control Probe (# 321831). Signal detection was achieved using HRP-based amplification with the following Opal dyes: Akoya Biosciences Opal™ 520 (Cat. No. FP1487001KT); Opal™ 570 (Cat. No. FP1488001KT), and Opal™ 690 (Cat. No. FP1497001KT). Sections were stained with DAPI and mounted using ProLong Gold antifade reagent (Invitrogen, Cat. No. P36930). Image registration of the HiPlex Assay was done with the RNAscope® HiPlex Image Registration Software (#300065).

### Immunohistochemistry

Mice were anesthetized with 5% isofluorane and perfused transcardially with 0.1 M saline phosphate buffer (PBS; [in mM] 154 NaCl, 13 Na2HPO4, 2.5 NaH2PO4, pH 7.4) followed by with 4% paraformaldehyde (PFA) in PBS. Spinal cords were then extracted by laminectomy, postfixed for 2 hours in same fixative, and cryoprotected in 30% sucrose / PBS solution at 4°C. Lumbar spinal cord sections were cut using a cryostat (Leica Microsystems). Transverse sections (25 μm-thick) were cut and placed in a 24-well plate containing PBS and stored at 4°C. Tissue sections were washed three times in PBS/0.3% Triton X-100 (PBS-T; Sigma, St. Louis, MO), and incubated in 10% normal donkey serum/PBS/0.3% Triton X-100 (NDST) for 1 hour, before adding the primary antibodies over 48 hours at 4°C in a 1% NDST solution. Sections were then washed three times with PBS-T solution and incubated for 1 hour with Alexa fluorophore-conjugated secondary antibodies diluted 1:500 in 1% NDST at room temperature (RT). The sections were washed three times with PBS, counterstained with NucBlue Fixed Cell Stain (DAPI) for 5 minutes at room temperature, mounted on slides, and coverslipped using Aqua Polymont (Polysciences, Inc., Warrington, USA). The slides were stored protected from light at 4°C until imaging.

Primary antibodies were used in 1% NDST at the indicated concentrations: anti-Homer1 rabbit (1:1000; Fronter Institute #MSFR103200), anti-VGluT1 guinea pig (1:1000, Millipore Sigma #AB5905), anti-gephyrin mouse (1:1000, Synaptic Systems #147021), anti-Pax2 rabbit (1:300, Life Technologies #716000), anti-GFP chicken (1:1000, Abcam #ab13970), anti-PV rabbit (1:1000, Swant #PV27), anti-Tlx3 rabbit (1:1000, Invitrogen #PA5-34555), anti-Cplx1 rabbit (1:1000, Abcam #ab231347), anti-HA mouse (1:1000, BioLegend #901516). All primary staining were detected using species-appropriate Alexa Fluor-conjugated secondary fluorophores.

### Microscopy and image analyses

Images for quantitative analysis were acquired using a Zeiss LSM780 scanning confocal microscope. All mouse/virus validation and rabies tracing imaging were collected using a 20x/0.40LD Plan Neofluar objective lens at 1024×1024 pixels/frame, 0.6X digital zoom, 12-bit depth, and detection using a 32 GaAsp detecter array. Synaptic marker imaging was performed on 63X Plan Apochromat NA=1.4 oil-immersion objective lens at 1024x1024 pixels/frame, 5X digital zoom, 12-bit depth, a z-step size of 0.44 μM, and the same detector array. The lasers were: laser diode 405nm 30mW, Ar Ion laser 458/488/514nm 25mW, DPSS-laser 561nm 20mW, HeNe RED 633nm 5mW. Image files were processed for analysis in ImageJ (NIH) using the “Blind Analysis Tools” plug-in to ensure experimenter blinding. High-resolution confocal stacks were first deconvolved using Huygen Essentials software using a full maximum likelihood extrapolation algorithm (Scientific Volume Imaging, Hilversum, The Netherlands). To quantify intensity of synaptic markers on PVNs and Cplx1-positive cells, three-dimensional surfaces of PVNs or Cplx1-positive cells were generated using the surface function in Imaris (Bitplane) was used to create neuronal masks. Signal from channels corresponding to synaptic proteins of interest was restricted to PVNs or Cplx1-positive cells using these masks to ensure puncta localization within neurons. The intensity and number of synaptic puncta localized to PVN somas and dendrites and on Cplx1-positive cells were then quantified manually.

### Chronic constriction injury surgery

Unilateral chronic constriction injury of the sciatic nerve (CCI) was done as previously described^64^. Three ligatures (6-0 Perma-Hand silk suture; Ethicon) were placed around the sciatic nerve bundle and loosely closed to construct but not arrest epineural blood flow. Mice developed robust mechanical allodynia that becomes chronic after two weeks and can persist for up to 8 weeks post-surgery.

### Viral injections – AAV and rabies

For the intraspinal injections, mice were anesthetized with 4% isoflurane/oxygen and maintained with 2% during the operation. A 1.5 cm incision was made on the back of the animal to expose the underlying spinal column at the lumbar level. Muscle and connective tissues overlaying L4-L5 spinal segments were removed, and the spinal column was immobilized in a stereotaxic frame. A glass microelectrode was lowered through the dura mater at the intervertebral space between the T13 and L1 vertebral segments (AP 0, ML 0.45, DV 0.25 mm) and 350 nL of virus was injected (50 nl/min). The electrode was slowly removed after 5 min. Carprofen analgesic (5 mg/kg) was given 45 mins prior to the beginning of the surgery and every 24 h for 3 days after the surgery. Mice recovered on a heat pad before being returned to their cage. Mice continued to be group housed for at least 2 weeks to allow for viral expression before behaviour or additional viruses were injected.

For the rabies tracing, 2 weeks after the initial AAV injection to allow the expression of the helper proteins, the surgical site was reopened, and the rabies virus was injected into the spinal cord at the same location. After waiting 5 days to allow for the rabies to infect axons and move retrograde transsynaptically, mice were euthanized for immunohistochemistry or fluorescent in situ hybridization.

For the intraperitoneal injection in neonatal mice, at postnatal day 5 (P5), mice were anesthetized by brief hypothermia on ice and administered a 15 µL intraperitoneal injection of AAV.

AAVs and their concentrations used in this study were as follows:

AAV2/php.S-CAG-ChR2-eGFP (2.0 ^E^13 GC/mL; Canadian Neurophotonics Platform [CNP] #4956);

AAV2/8-hSyn-FLEX-TVA-P2A-eGFP-2A-oG (1.0 ^E^13 GC/mL; CNP #5446);

RABV-EnvA-deltaG-mCherry (1.0^E^9 TU/mL; CNP #3.8);

AAV2/8-hSyn1-hSyn1-roxSTOP-dlox-TVA_2A(TAV)_RabG(rev)-WPRE-dlox-hGHp(A) (1.5 ^E^12 vg/mL from lab of Dr. Zeilhofer at University of Zurich);

AAV2/1-hEF1a-hTLV1-Con/Don(hM4D(Gi)_mCherry)-WPRE-hGHp(A) (5.3 ^E^12 vg/mL from lab of Dr. Zeilhofer at University of Zurich);

AAV2/1-hEF1a-hTLV1-Con/Don(HA_hM3D(Gq))-WPRE-hGHp(A) (4.8 ^E^12 vg/mL from lab of Dr. Zeilhofer at University of Zurich).

### Electrophysiology

Transverse lumbar spinal cord slices and coronal brain slices were obtained from adult mice (6-12 weeks old). Animals were deeply anesthetized through an intraperitoneal injection of 2,2,2-Tribromoethanol (Avertin, 250mg/kg). The spinal cord was quickly removed and transferred into an ice-cold (4°C) oxygenated (95% O2, 5% CO2) N-Methyl-D-Glucamine based artificial cerebrospinal fluid (NMDG-ACSF) solution (bubbled with 95% O2 and 5% CO2) containing the following (in mM): 93 NMDG, 2.5 KCl, 1.25 NaH2PO4, 30 NaHCO3, 20 HEPES, 25 glucose, 2 thiourea, 5 Na-L-ascorbate, 3 Na-pyruvate, 12 N-acetyl-L-cysteine, 0.5 CaCl2/2H2O, and 10 MgSO4/7H2O (pH 7.3–7.4 adjusted with HCl 12M). After removing the dura mater and ventral roots, the spinal cord was embedded into 3% low-melting agarose (Fisher Scientific BP165-25) and serial transverse slices (530-570 µm) were cut from the lumbar spinal cord with the dorsal roots and DRGs still attached using a vibratome (Leica VT1200S). Slices were incubated at 36°C containing HEPES-based recovery ACSF for 30 minutes, equilibrated with 95% O2 and 5% CO2. Following the recovery incubation, slices were transferred to a recording chamber and continuously superfused with oxygenated ACSF (in mM): 119 NaCl, 24 NaHCO3, 2.5 KCl, 1.25 NaH2PO4, 2 CaCl2, 2 MgCl2, and 12.5 glucose (bubbled with 95% O2 and 5% CO2; pH 7.3; 300±5 mOsm measured), where they were then maintained at room temperature prior to transfer to the recording chamber. Slices were transferred to a recording chamber (volume approximately 1 ml) and held down with a pewter wire. All recordings were performed at room temperature.

Slices were superfused continuously with oxygenated ACSF (2 mL/min). Patch pipettes were pulled from borosilicate glass capillaries (Harvard Apparatus) with a P-97 puller (Sutter Instruments). They were filled with a solution containing (in mM) 135 K-Gluconate, 6 NaCl, 2 MgCl2, 10 HEPES, 0.1 EGTA, 2 MgATP, 0.8 NaGTP (pH 7.3-7.4, adjusted with KOH; osmolarity, 300 mOsm, adjusted with sucrose) and had final tip resistances of 7–9 MΩ for whole-cell recording. For the mIPSCs recordings, a cesium-chloride based internal solution was used containing (in mM): 130 CsCl, 10 HEPES, 10 EGTA, 1 MgCl2, 2 MgATP, 0.3 NaGTP (pH 7.3-7.4, adjusted with CsOH; osmolarity, 300 mOsm, adjusted with sucrose) ^65^. Neurons were viewed by an upright microscope (Olympus) with a 40X water-immersion objective, infrared differential interference contrast (IR-DIC) and fluorescence. Whole-cell patch clamp recordings were made in identified PVNs expressing tdTomato and acquired with pClamp 10.0 software (Molecular Devices) using MultiClamp 700B patch clamp amplifier and Digidata 1440A (Molecular Devices). Recordings were low pass filtered on-line at 2 kHz, digitized at 20 kHz and stored on a PC using pClamp software (Molecular Devices). Series resistance was monitored throughout the experiments and was not compensated. Data were discarded if series resistance varied more than ±20 MΩ. Voltage clamp data were recorded at a holding potential of-70 mV for eEPSCs/oEPSCs and-40mV for eIPSCs/oEPSCs.

Dorsal roots were stimulated 5 times per protocol using a suction electrode at 25, 100, and 500 µA to activate Aβ, or Aβ + Aδ, or Aβ + Aδ + C fibers, respectively, at varying frequencies of 0.05 Hz, 0.1Hz, 1Hz, 2Hz, 4Hz, 20Hz, 40Hz (duration 0.1 ms) using a A365 stimulus isolator (World Precision Instrument) as previously described ^28^. To address the monosynaptic nature of the Aβ fiber evoked response, the root was systematically stimulated five times at 5-75 µA (+10 µA increments) and 2Hz. EPSCs were considered monosynaptic in an absence of synaptic failure or latency >2ms. Conduction velocity was measured and estimated based on the length of the root from the tip of the suction electrode to the recorded PVN. Intrinsic excitability of PVNs receiving dorsal root input was assessed using two current-clamp protocols: (1) a 1-second depolarizing current step of 150 pA, and (2) step current injections delivered at 20 Hz in 10 pA increments from 0-80 pA.

To measure mEPSCs of PVNs, the K-gluconate based internal solution was used, with bath application of 1 µM TTX (Cedarlane #14964-1), 15 µM bicuculline (Alomone Labs #B-137), and 500 nM strychnine (Sigma Millipore #S8753). To measure mIPSCs of PVNs, the CsCl based internal solution was used, with bath application of 1 µM TTX and 10 µM NBQX (Alomone Labs #N-186). To measure sEPSCs, the same respective internal solutions were used in the absence of bath-applied pharmacological agents.

To investigate the inhibitory nature on PVNs, three pharmacological protocols were used: (1) bath application of 15 µM bicuculline and 500 nM strychnine, (2) bath application of 10 µM NBQX, and (3) bath application of 1 µM TTX and 200 µM 4-AP (Millipore Sigma #A78403). Drugs were bath applied for at least 5 mins prior to post-drug recordings. For all protocols, primary afferent inputs were activated using either electrical or optogenetic stimulation. The eEPSCs/oEPSCs and eIPSCs/oIPSCs were recorded before and after drug application, as described above. Optogenetic stimulation of primary afferents was delivered using the same stimulation frequencies as electrical dorsal root stimulation, with 5 ms full-field blue light pulses (460nm, 5.50 mW) generated by an LED light source (CoolLED pE-300 Ultra). Light was stimulated on the entire spinal dorsal horn.

To examine PVN-PVN feedforward inhibition from Aβ input, both dorsal root electrical stimulation and photo-stimulation of the spinal doesal horn with the blue LED light were used. Only PVNs that responded with IPSP in current clamp were used. PVNs were held at 0 mV as only electrically (0.1 ms duration) and optically (0.5 ms, depletion stimulations: 90 ms, 30ms, 450 ms duration) evoked inhibitory current was measured.

### Electrophysiology Analyses and Clustering

Action potential properties and intrinsic excitability analyses were performed on MATLAB software with the toolbox, ElecFeX ^66^. For the analyses of the synaptically evoked responses of PVNs, electrophysiological traces were downsampled to 10kHz prior to clustering. Each PVN was stimulated five times, with each individual response treated independently to capture trial-to-trial variability (one response per row). Responses from all 7 experimental conditions were pooled into a single data matrix, and z-scored for subsequent analysis. Principal component analysis (PCA) was performed in MATLAB (R2024a) to reduce dimensionality. Clustering was performed using a Gaussian mixture model (GMM) applied to the first four principal components, which together accounted for 82% of the total variance. The optimal number of clusters was determined using the Bayesian information criterion (BIC). Each PVN response was evaluated for their probability to belong to each gaussian and finally assigned to the cluster with the highest probability. Uniform manifold approximation and projections (UMAPs) computed using default parameters were used for low-dimensional visualization of synaptically evoked responses of PVNs across experimental conditions.

### Behaviour

All mice were habituated to the experimenter handling and behaviour room the day before testing began. Experimenter was blind to the experimental groups. Male and female mice were tested separately. Mice would receive an intraperitoneal injection of either deschloroclozapine dihydrochloride (DCZ, 3μg/kg, HelloBio, #HB9126) or sterile saline. Testing would take place between 30 mins to 90mins post-injection.

### von Frey assay

Static mechanical sensitivity was assessed as previously described ^25^. Mechanical nocifensive threshold was assessed by placing mice on an elevated wire-mesh grid and stimulating the plantar surface of the hindpaw with von Frey filaments. Starting with a low weight filament, each filament was applied 5 times against the hindpaw. If no nocifensive response was elicited after the 5 applications, the next filament (stronger) was applied and so on until 5 of 5 applications elicited a response. The filament where 5 of 5 applications elicited a nocifensive response (licking, biting, paw guarding, flicking, escape behaviours) was deemed the mechanical threshold. Before slice electrophysiology of CCI mice, we verified their development of mechanical allodynia by testing mechanical sensitivity before and post CCI.

### Dynamic brush assay

A fine, modified paintbrush where the bristles are plucked off with only a handful of fine hairs remaining is used to swept across the center of the hindpaw, from heel to toes 5 times. The mechanical behaviour response of the mice to the brushes was recorded as no response, paw withdrawal, or nocifensive reaction.

### Radiant heat paw-withdrawal assay

The withdrawal latency to high temperature was measured using the radiant heat paw-withdrawal (Hargreaves) test (Stoelting). Mice were placed into individual restrainers on a glass platform 60 min before the experiment and allowed to habituate. A noxious thermal stimulus was focused through the glass onto the plantar surface of a hindpaw until the animal withdrew the paw from the heat source. The paw-withdrawal latency was automatically measured to the nearest 0.1 s. A cut-off latency of 20 s was used to avoid tissue damage.

### Acetone cold sensitivity assay

A droplet of acetone (Sigma-Aldrich, #179124) is applied to the plantar surface of the hind paw. The paw is observed for the duration of the nocifensive response for up to 60 seconds.

### Graphs and statistics

All graphs and statistics were generated with either Graphpad Prism 10 or MATLAB R2024a. Values were reported as mean ± standard error of the mean (SEM).

## RESOURCE AVAILABILITY

### Lead Contact

Requests for further information and resources should be directed to and will be fulfilled by the lead contact, Reza Sharf-Naeini.

### Materials availability

This study did not generate new materials.

## ACKNOWLEDGMENTS

We thank Dr. Arjun Krishnaswamy and his lab members, Dr. Aline Giselle Rangel Olguin and Mr. Rian Fritz Jalandoni, for their help with the clustering analyses. We thank the McGill Animal Behavioural Characterization platform for the mouse behaviour experimental support.

H.Q. was funded by the Canadian Institute of Health Research (CIHR) Doctoral award. R.S.-N. was supported by a project grant from the CIHR (PJT-162404).

## Author contributions

Conceptualization: HQ, RSN

Methodology: HQ

Visualization: HQ

Funding acquisition: RSN

Writing – original draft: HQ

Writing – review & editing: HQ, CP, HUZ, RD, RSN

## Declaration of interests

The authors declare no competing interests.

## Classification

Biological Sciences; Neuroscience

## Notes

### Competing Interest Statement

The authors have declared no competing interest.

