## Supplementary Information for "Peripheral nerve injury reallocates primary afferent input through spinal parvalbumin microcircuits"

Mailing address:

3649 Promenade Sir William Osler

Bellini Life Sciences Center, McGill University

Montreal, Quebec, Canada H3G 1Y7

**This PDF file includes:**

Supplementary Figures 1 to 14

Supplementary Table 1

### Figure S1

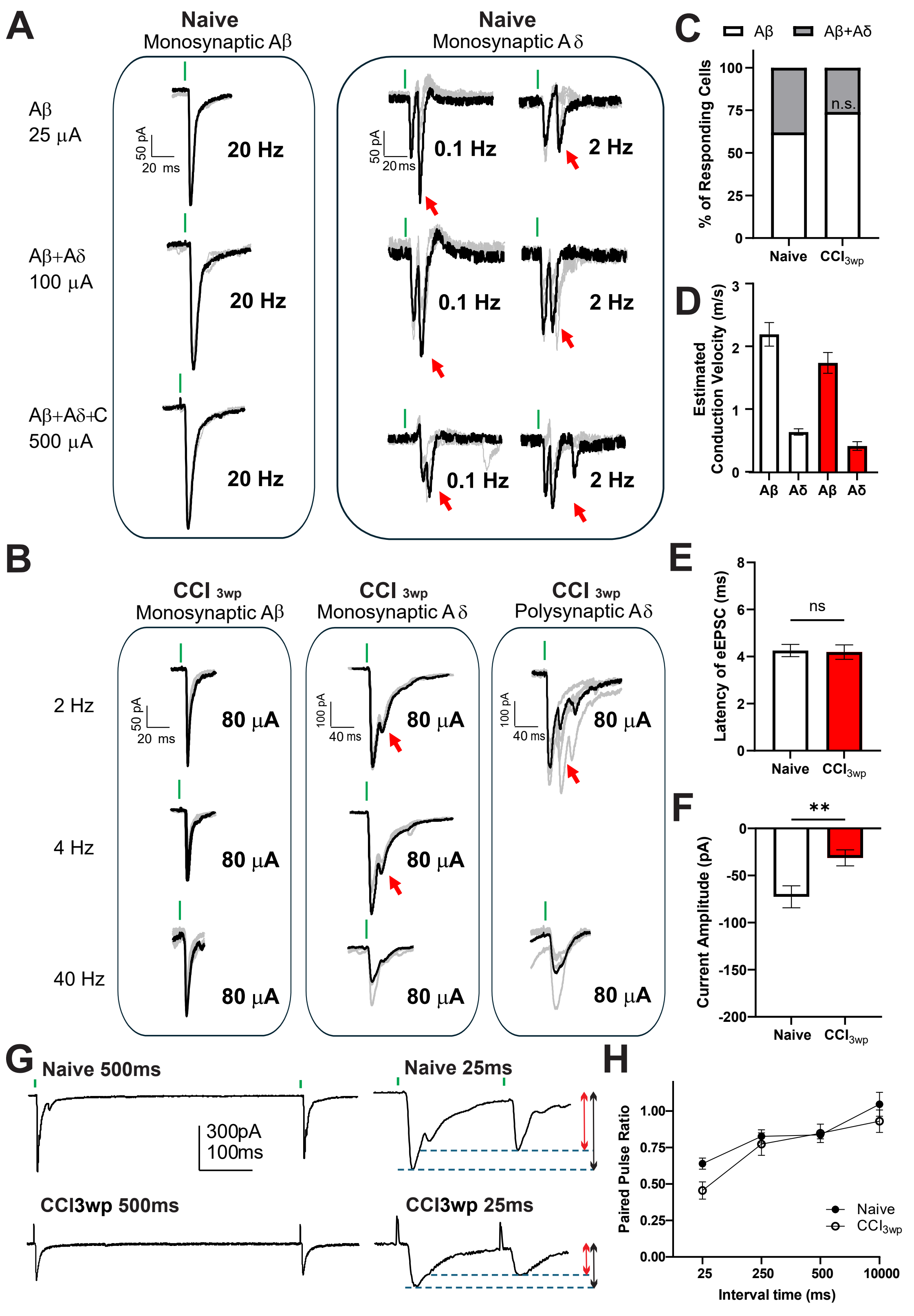
A
Naive
Monosynaptic Aβ
Naive
Monosynaptic Aδ
Aβ
25 μA
Aβ+Aδ
100 μA
Aβ+Aδ+C
500 μA
50 pA
20 ms
20 Hz
0.1 Hz
2 Hz
C
Aβ
Aβ+Aδ
% of Responding Cells
n.s.
Naive
CCI3wp
D
Estimated Conduction Velocity (m/s)
Aβ
Aδ
B
CCI 3wp
Monosynaptic Aβ
Monosynaptic Aδ
Polysynaptic Aδ
2 Hz
4 Hz
40 Hz
80 μA
100 pA
40 ms
E
Latency of eEPSC (ms)
ns
F
Current Amplitude (pA)
**
G
Naive 500ms
Naive 25ms
300pA
100ms
CCI3wp 500ms
CCI3wp 25ms
H
Paired Pulse Ratio
Interval time (ms)
Naive
CCI3wp

### Figure S2

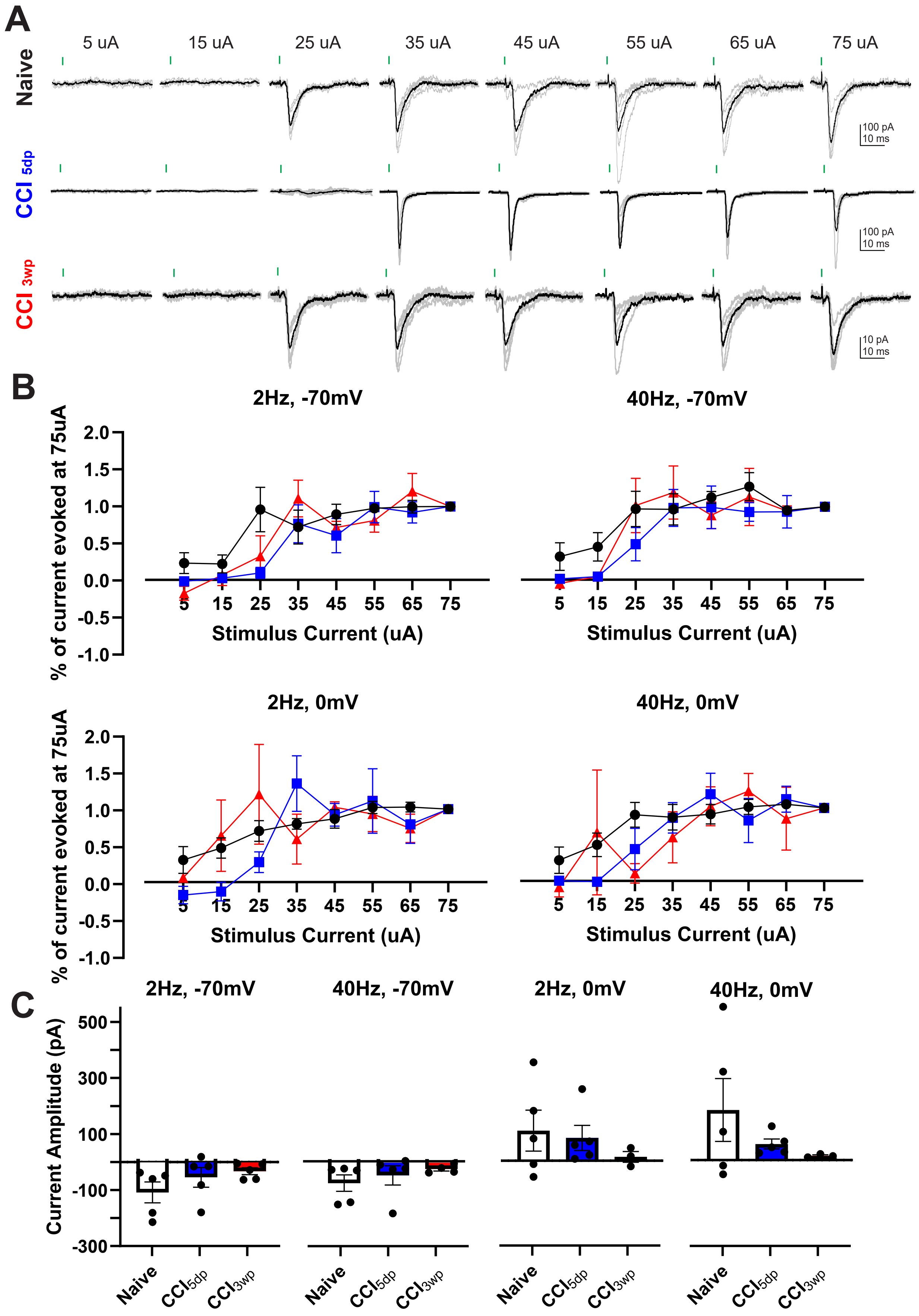

A
5 uA
15 uA
25 uA
35 uA
45 uA
55 uA
65 uA
75 uA
Naive
CCI 5dp
CCI 3wp
100 pA
10 ms
100 pA
10 ms
10 pA
10 ms
B
2Hz, -70mV
40Hz, -70mV
% of current evoked at 75uA
Stimulus Current (uA)
2Hz, 0mV
40Hz, 0mV
% of current evoked at 75uA
Stimulus Current (uA)
C
2Hz, -70mV
40Hz, -70mV
2Hz, 0mV
40Hz, 0mV
Current Amplitude (pA)
Naive
CCI5dp
CCI3wp

### Figure S3

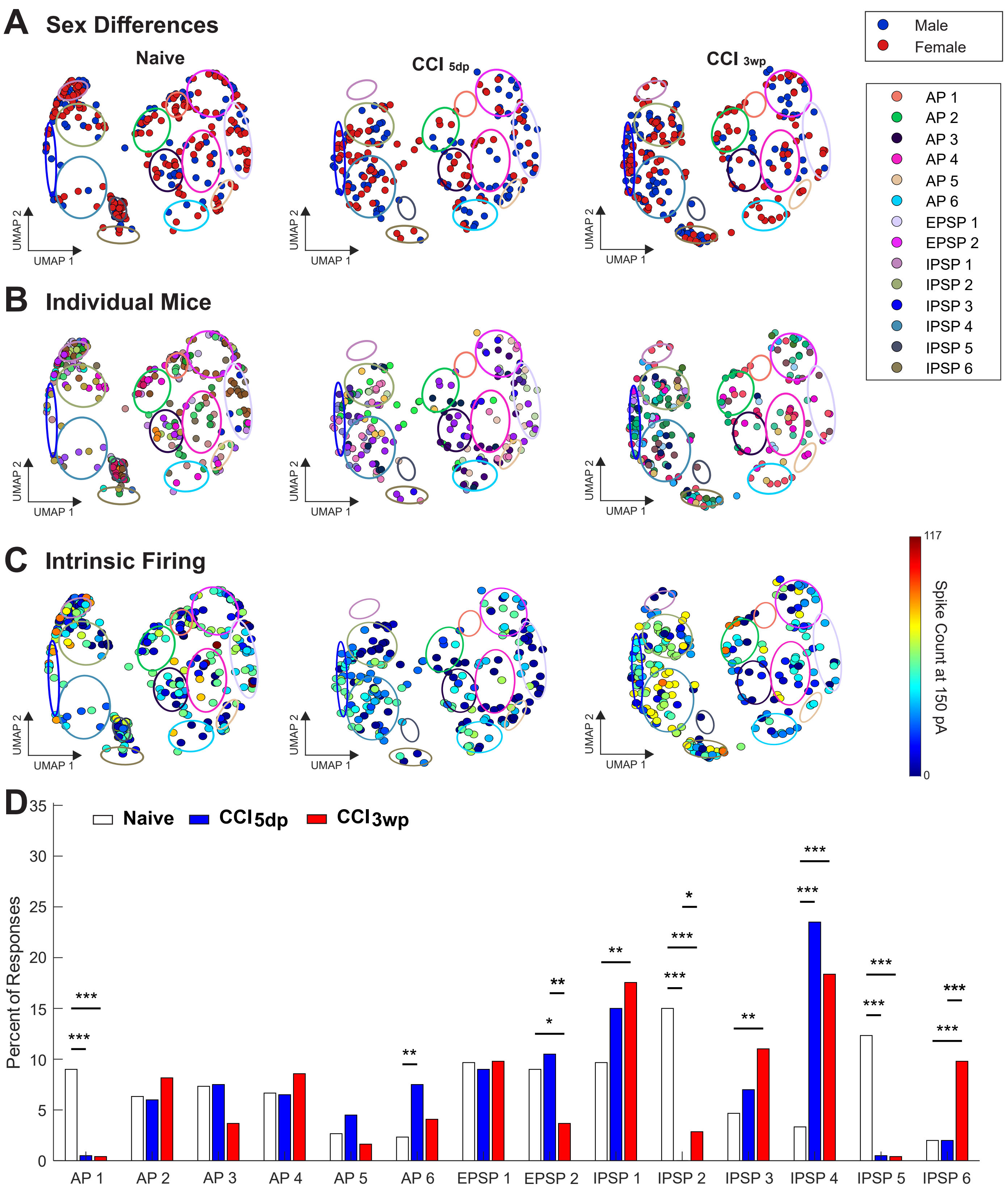
A Sex Differences
Naive
CCI 5dp
CCI 3wp
Male
Female
AP 1
AP 2
AP 3
AP 4
AP 5
AP 6
EPSP 1
EPSP 2
IPSP 1
IPSP 2
IPSP 3
IPSP 4
IPSP 5
IPSP 6
UMAP 1
UMAP 2
B Individual Mice
C Intrinsic Firing
Spike Count at 150 pA
117
0
D
Naive
CCI5dp
CCI3wp
Percent of Responses
0
5
10
15
20
25
30
35
AP 1
AP 2
AP 3
AP 4
AP 5
AP 6
EPSP 1
EPSP 2
IPSP 1
IPSP 2
IPSP 3
IPSP 4
IPSP 5
IPSP 6

### Figure S4

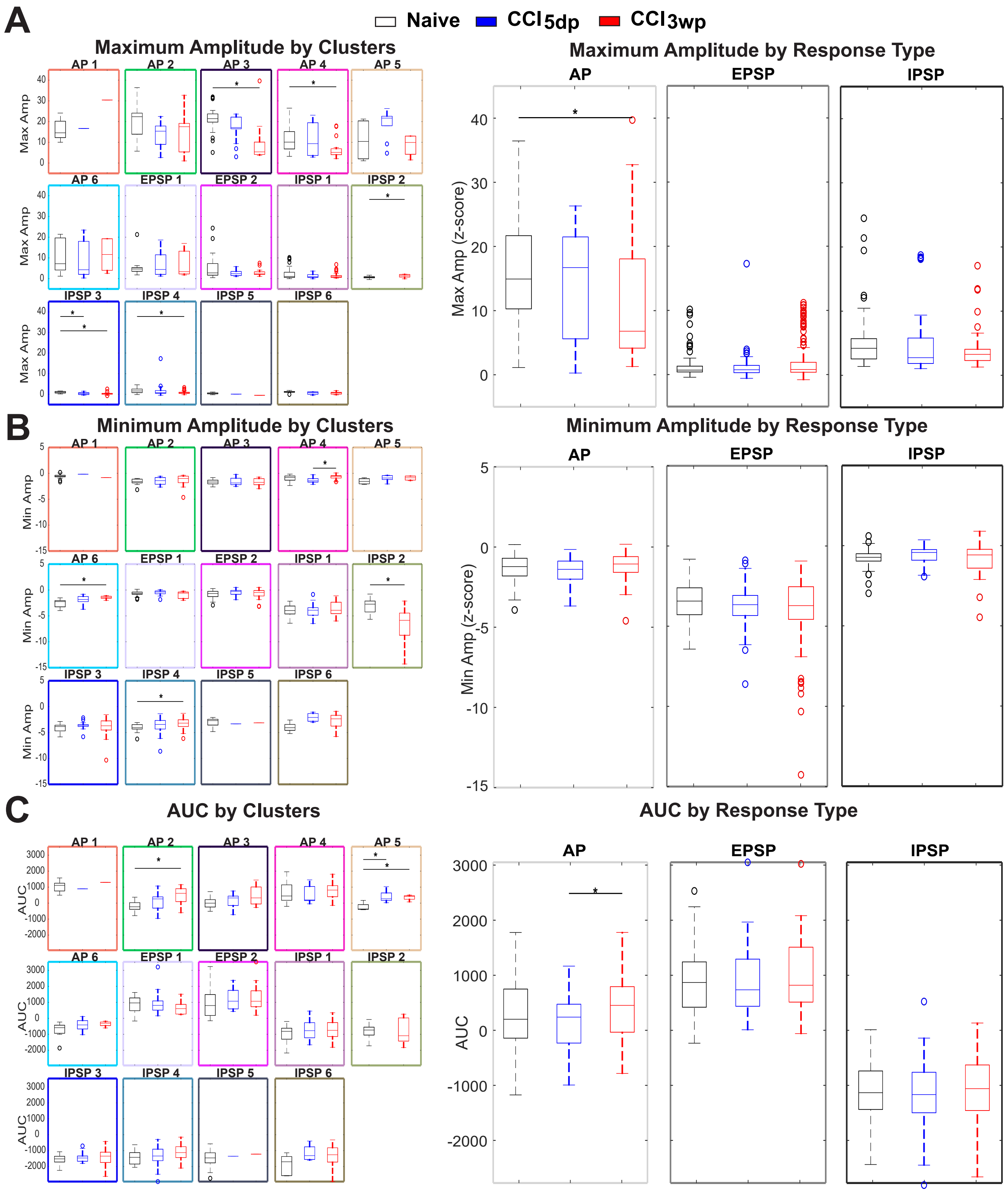
A
B
C
Naive
CCI5dp
CCI3wp
Maximum Amplitude by Clusters
Maximum Amplitude by Response Type
Minimum Amplitude by Clusters
Minimum Amplitude by Response Type
AUC by Clusters
AUC by Response Type
AP 1
AP 2
AP 3
AP 4
AP 5
AP 6
EPSP 1
EPSP 2
IPSP 1
IPSP 2
IPSP 3
IPSP 4
IPSP 5
IPSP 6
AP
EPSP
IPSP
Max Amp
Max Amp (z-score)
Min Amp
Min Amp (z-score)
AUC

### Figure S5

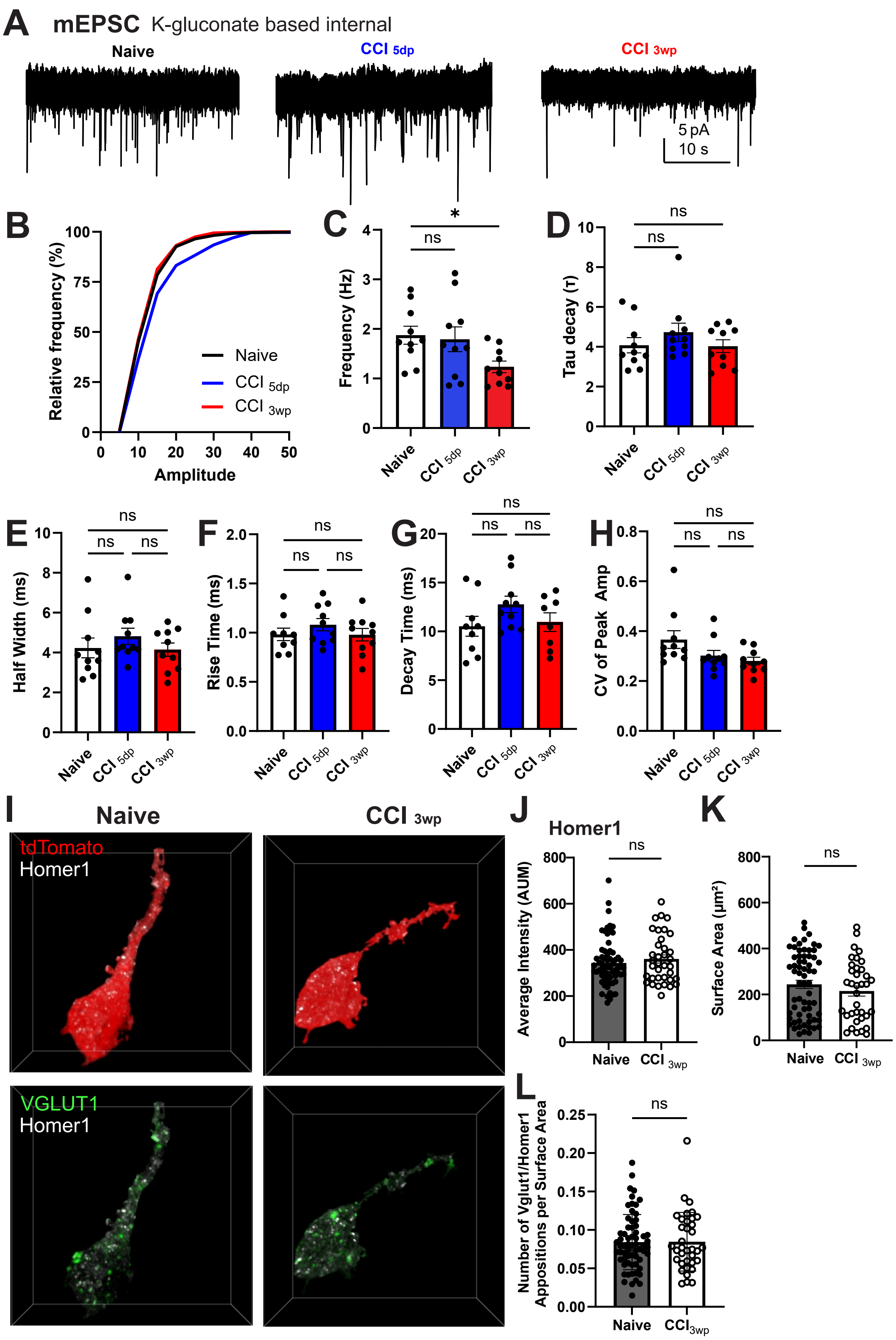
A
mEPSC K-gluconate based internal
Naive
CCI 5dp
CCI 3wp
5 pA
10 s
B
Relative frequency (%)
Amplitude
Naive
CCI 5dp
CCI 3wp
C
Frequency (Hz)
D
Tau decay (τ)
E
Half Width (ms)
F
Rise Time (ms)
G
Decay Time (ms)
H
CV of Peak Amp
ns
*
Naive
CCI 5dp
CCI 3wp
I
Naive
CCI 3wp
tdTomato
Homer1
VGLUT1
Homer1
J
Homer1
Average Intensity (AUM)
K
Surface Area (µm²)
L
Number of Vglut1/Homer1 Appositions per Surface Area
Naive
CCI 3wp

### Figure S6

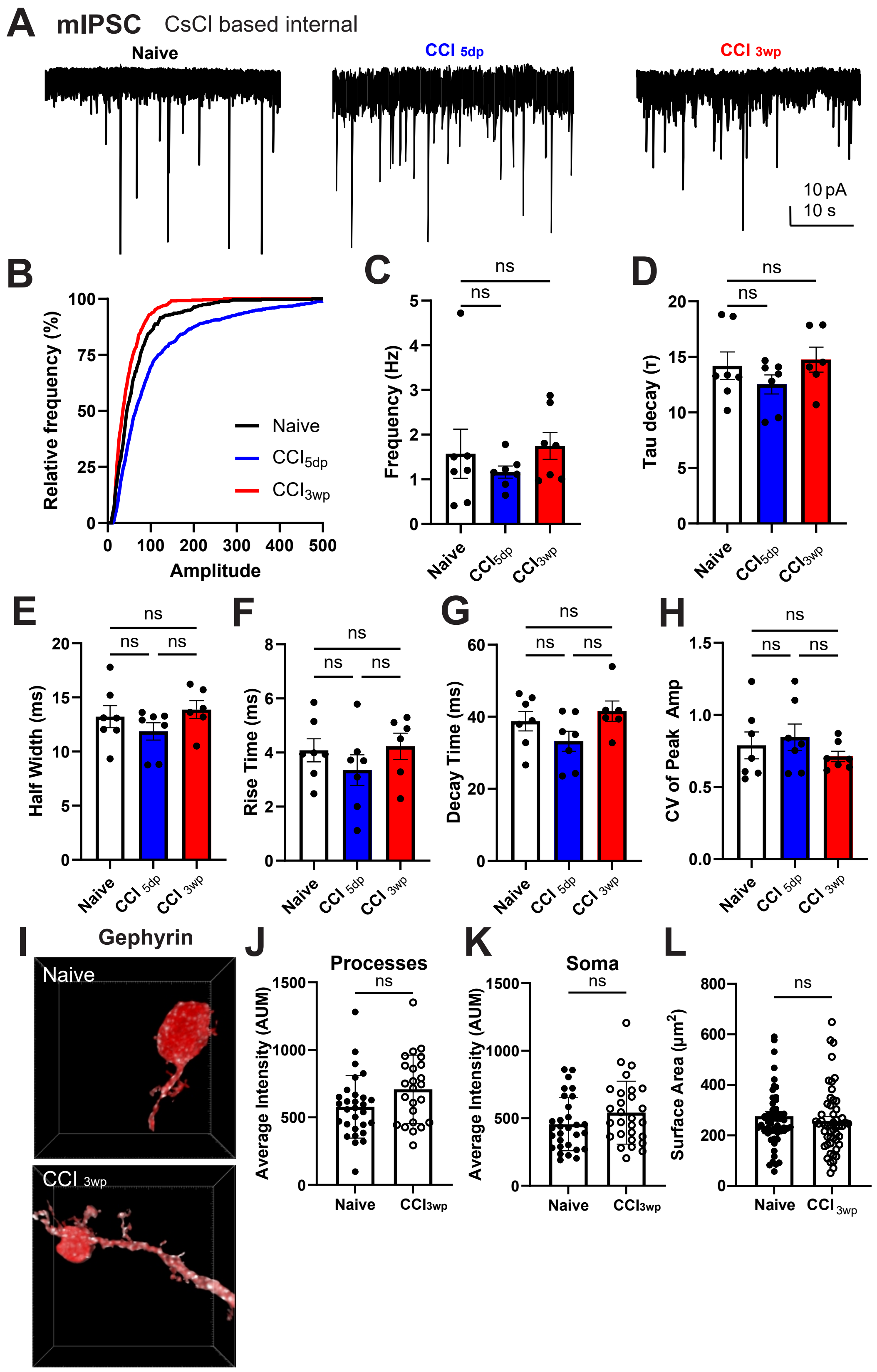
A
mIPSC
CsCl based internal
Naive
CCI 5dp
CCI 3wp
10 pA
10 s
B
Relative frequency (%)
Amplitude
Naive
CCI5dp
CCI3wp
C
Frequency (Hz)
ns
D
Tau decay (τ)
E
Half Width (ms)
F
Rise Time (ms)
G
Decay Time (ms)
H
CV of Peak Amp
I
Gephyrin
Naive
CCI 3wp
J
Processes
Average Intensity (AUM)
K
Soma
L
Surface Area (μm2)

### Figure S7

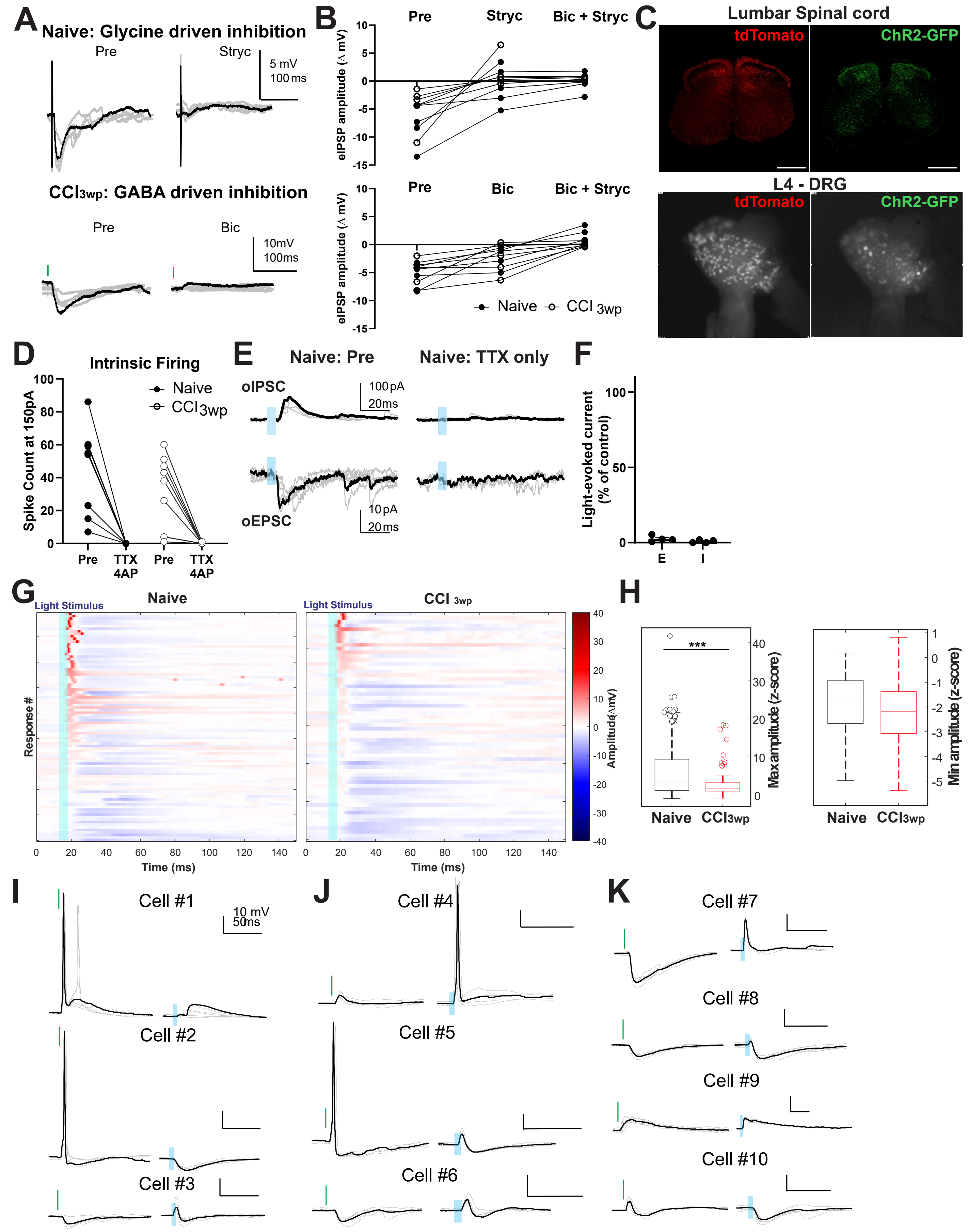


**Naive**

| Cell # | DR | ChR2 |
|---|---|---|
| 1 | AP | EPSP |
| 2 | AP | IPSP |
| 3 | IPSP | EPSP+IPSP |

**CCI 5dp**

| Cell # | DR | ChR2 |
|---|---|---|
| 4 | EPSP | AP |
| 5 | AP + IPSP | EPSP+IPSP |
| 6 | IPSP | EPSP+IPSP |

**CCI 3wp**

| Cell # | DR | ChR2 |
|---|---|---|
| 7 | IPSP | EPSP |
| 8 | IPSP | EPSP+IPSP |
| 9 | EPSP | EPSP |
| 10 | EPSP + IPSP | IPSP |

### Figure S8

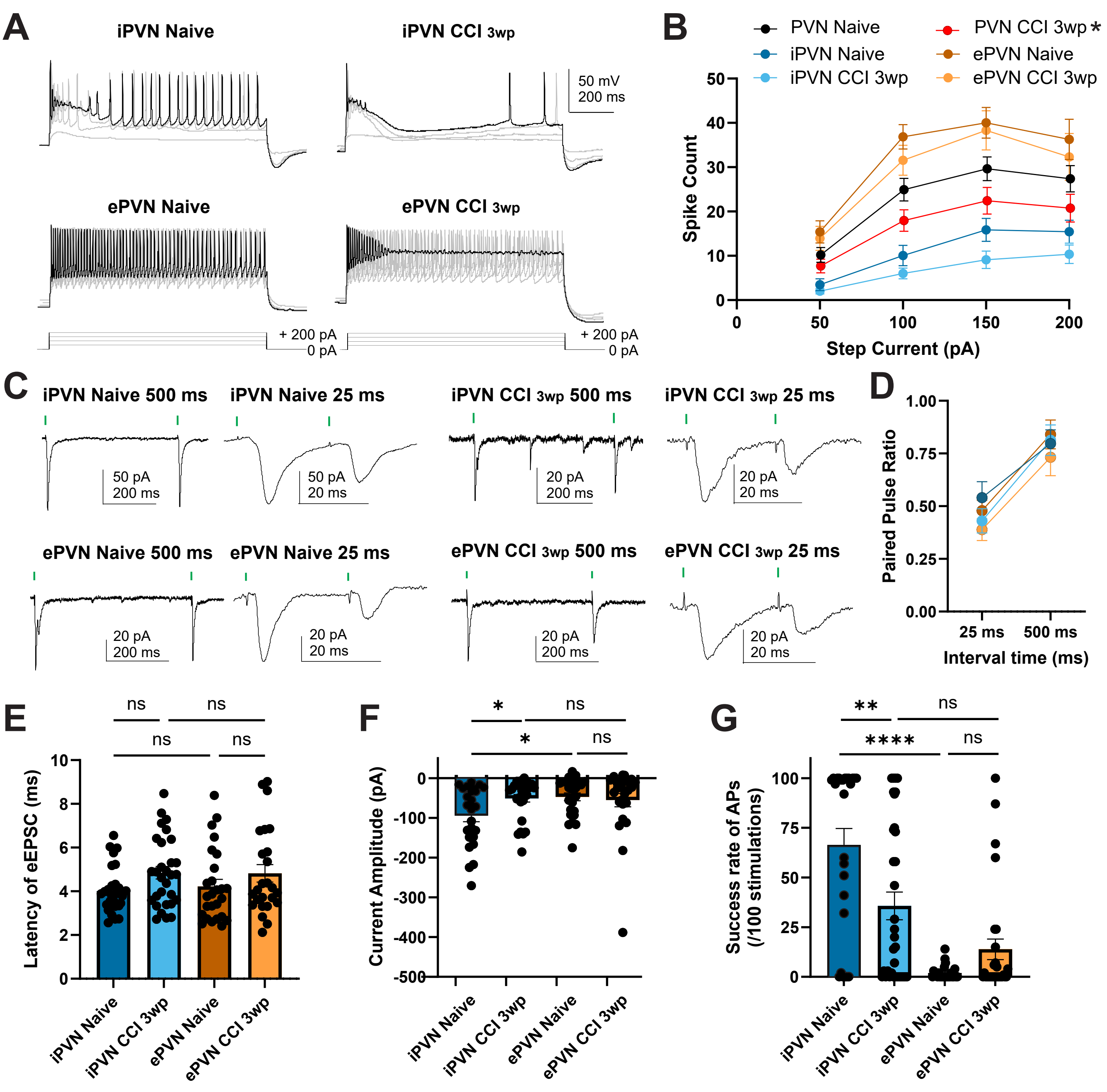

A
iPVN Naive
iPVN CCI 3wp
ePVN Naive
ePVN CCI 3wp
50 mV
200 ms
+ 200 pA
0 pA
B
PVN Naive
iPVN Naive
iPVN CCI 3wp
PVN CCI 3wp *
ePVN Naive
ePVN CCI 3wp
Spike Count
Step Current (pA)
C
iPVN Naive 500 ms
iPVN Naive 25 ms
iPVN CCI 3wp 500 ms
iPVN CCI 3wp 25 ms
ePVN Naive 500 ms
ePVN Naive 25 ms
ePVN CCI 3wp 500 ms
ePVN CCI 3wp 25 ms
50 pA
200 ms
20 pA
20 ms
D
Paired Pulse Ratio
25 ms
500 ms
Interval time (ms)
E
Latency of eEPSC (ms)
ns
F
Current Amplitude (pA)
*
ns
G
Success rate of APs (/100 stimulations)
**
****
ns
iPVN Naive
iPVN CCI 3wp
ePVN Naive
ePVN CCI 3wp

### Figure S9

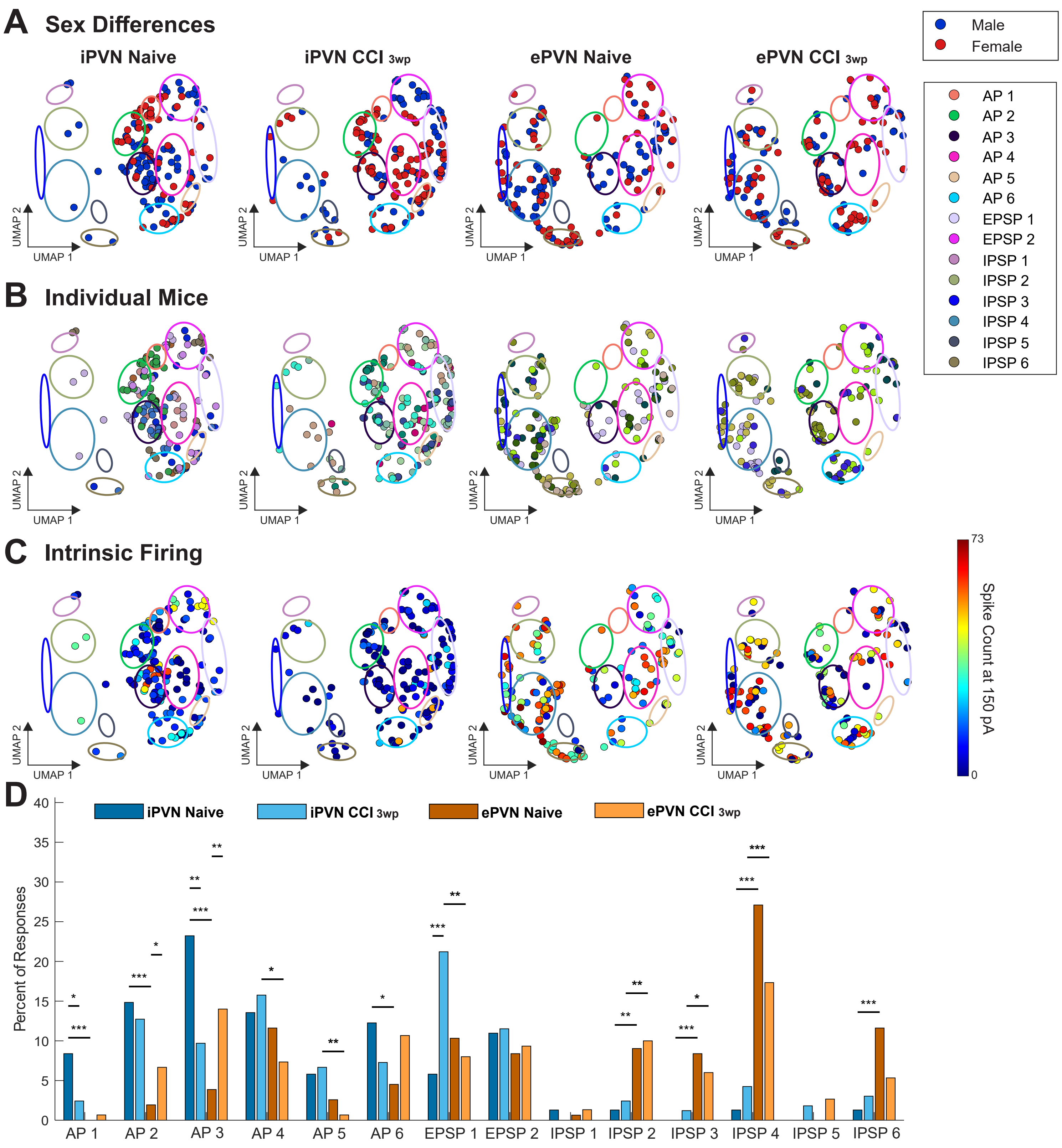

A Sex Differences
iPVN Naive
iPVN CCI 3wp
ePVN Naive
ePVN CCI 3wp
Male
Female
AP 1
AP 2
AP 3
AP 4
AP 5
AP 6
EPSP 1
EPSP 2
IPSP 1
IPSP 2
IPSP 3
IPSP 4
IPSP 5
IPSP 6
UMAP 1
UMAP 2
B Individual Mice
C Intrinsic Firing
Spike Count at 150 pA
73
0
D
Percent of Responses
iPVN Naive
iPVN CCI 3wp
ePVN Naive
ePVN CCI 3wp
0
5
10
15
20
25
30
35
40
AP 1
AP 2
AP 3
AP 4
AP 5
AP 6
EPSP 1
EPSP 2
IPSP 1
IPSP 2
IPSP 3
IPSP 4
IPSP 5
IPSP 6

### Figure S10

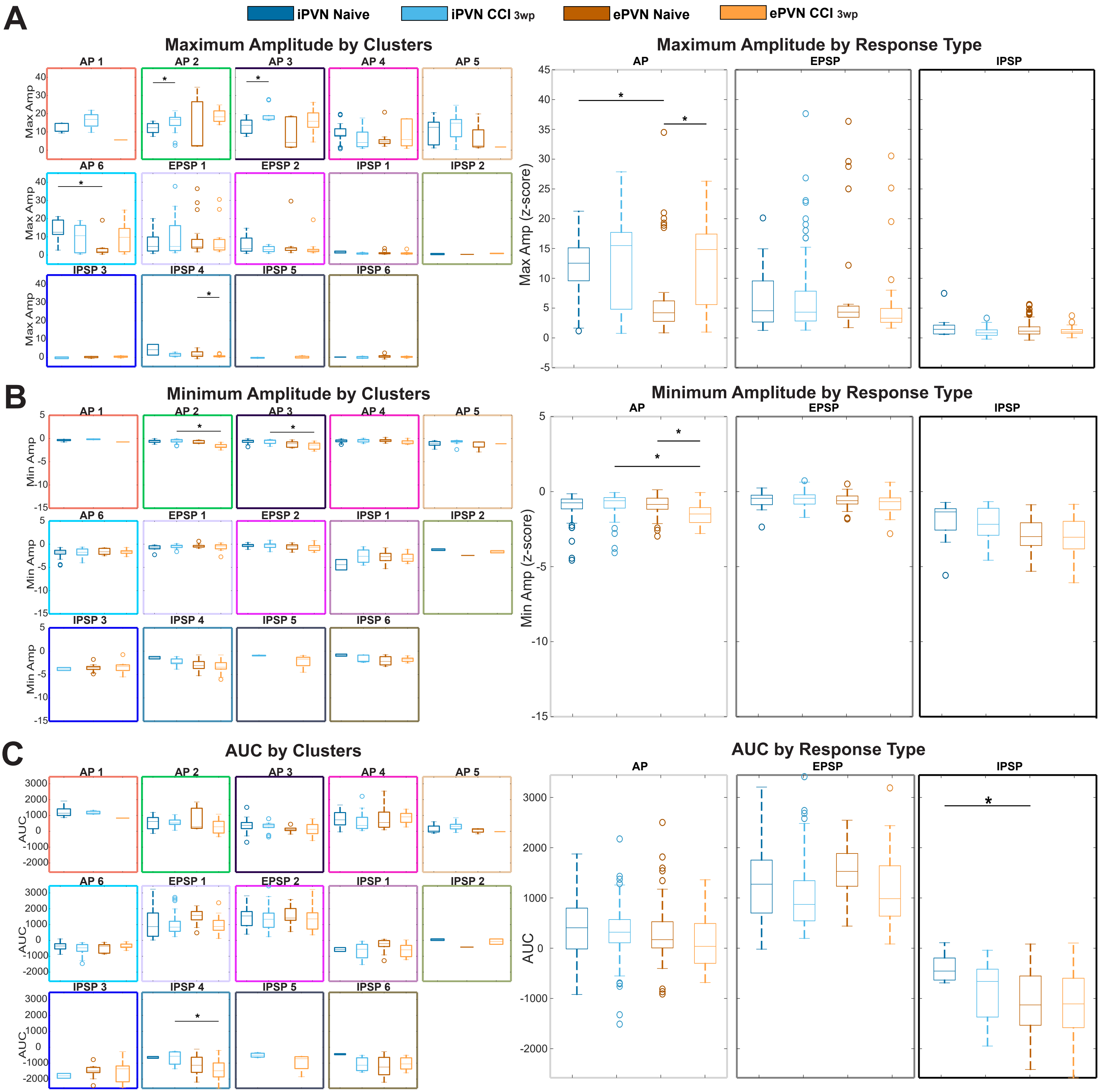
A
B
C
iPVN Naive
iPVN CCI 3wp
ePVN Naive
ePVN CCI 3wp
Maximum Amplitude by Clusters
Maximum Amplitude by Response Type
Minimum Amplitude by Clusters
Minimum Amplitude by Response Type
AUC by Clusters
AUC by Response Type
AP 1
AP 2
AP 3
AP 4
AP 5
AP 6
EPSP 1
EPSP 2
IPSP 1
IPSP 2
IPSP 3
IPSP 4
IPSP 5
IPSP 6
AP
EPSP
IPSP
Max Amp
Max Amp (z-score)
Min Amp
Min Amp (z-score)
AUC

### Figure S11

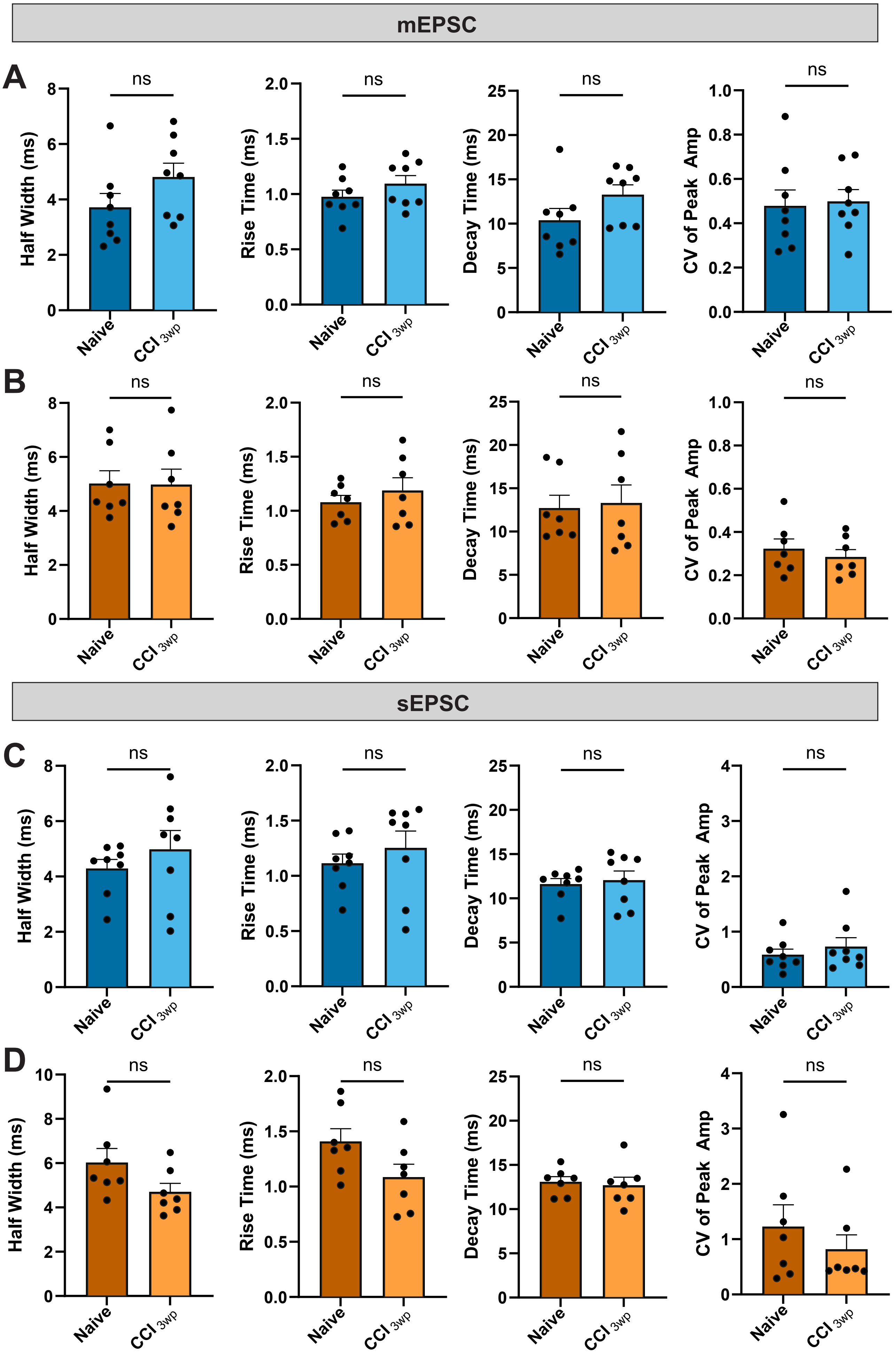
mEPSC
A
B
sEPSC
C
D
ns
Half Width (ms)
Rise Time (ms)
Decay Time (ms)
CV of Peak Amp
Naive
CCI 3wp

### Figure S12

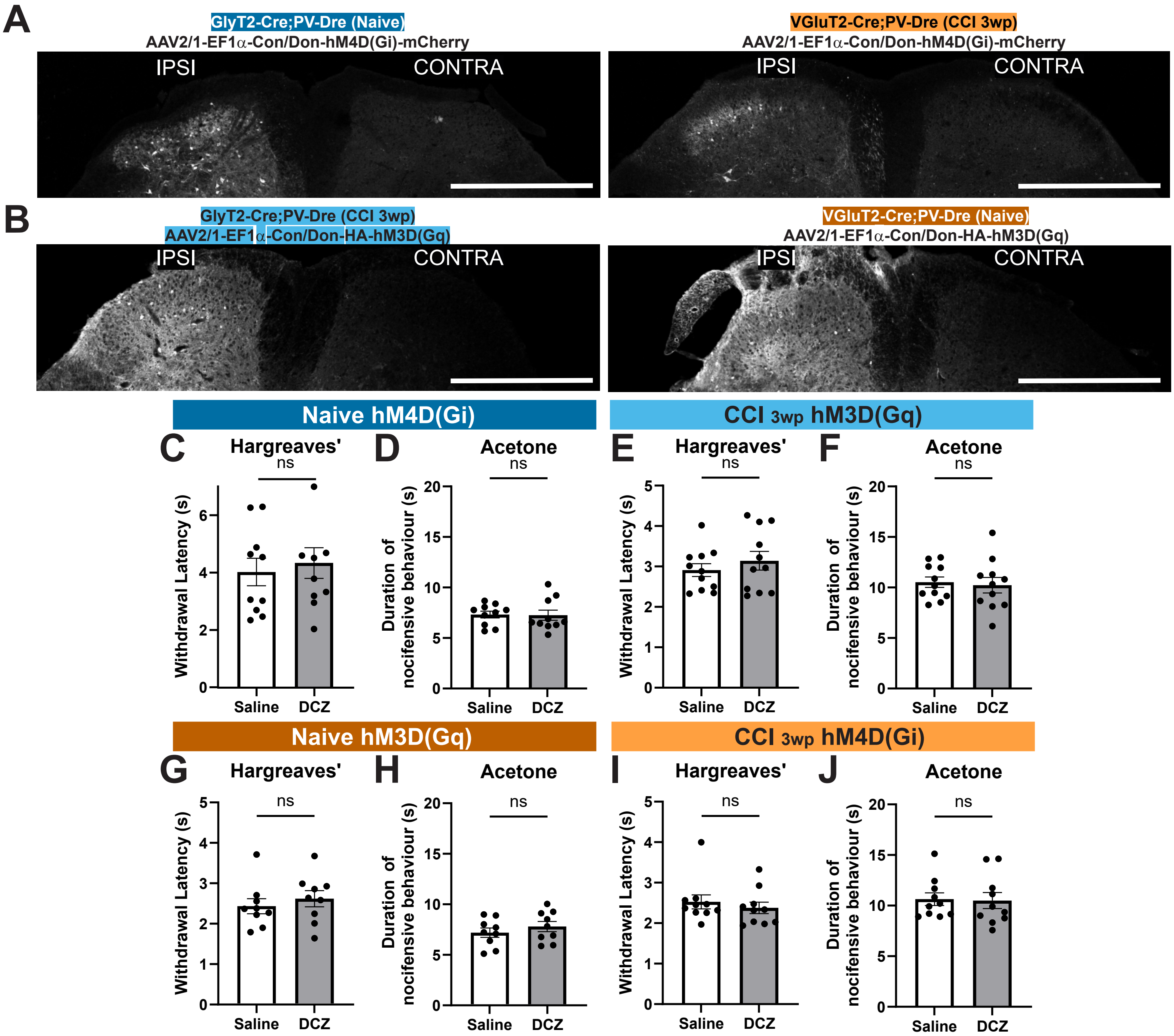

A
GlyT2-Cre;PV-Dre (Naive)
AAV2/1-EF1α-Con/Don-hM4D(Gi)-mCherry
IPSI
CONTRA
VGluT2-Cre;PV-Dre (CCI 3wp)
AAV2/1-EF1α-Con/Don-hM4D(Gi)-mCherry
B
GlyT2-Cre;PV-Dre (CCI 3wp)
AAV2/1-EF1α-Con/Don-HA-hM3D(Gq)
VGluT2-Cre;PV-Dre (Naive)
AAV2/1-EF1α-Con/Don-HA-hM3D(Gq)
Naive hM4D(Gi)
CCI 3wp hM3D(Gq)
C Hargreaves'
D Acetone
E Hargreaves'
F Acetone
ns
Withdrawal Latency (s)
Duration of nocifensive behaviour (s)
Saline
DCZ
Naive hM3D(Gq)
CCI 3wp hM4D(Gi)
G Hargreaves'
H Acetone
I Hargreaves'
J Acetone

### Figure S13

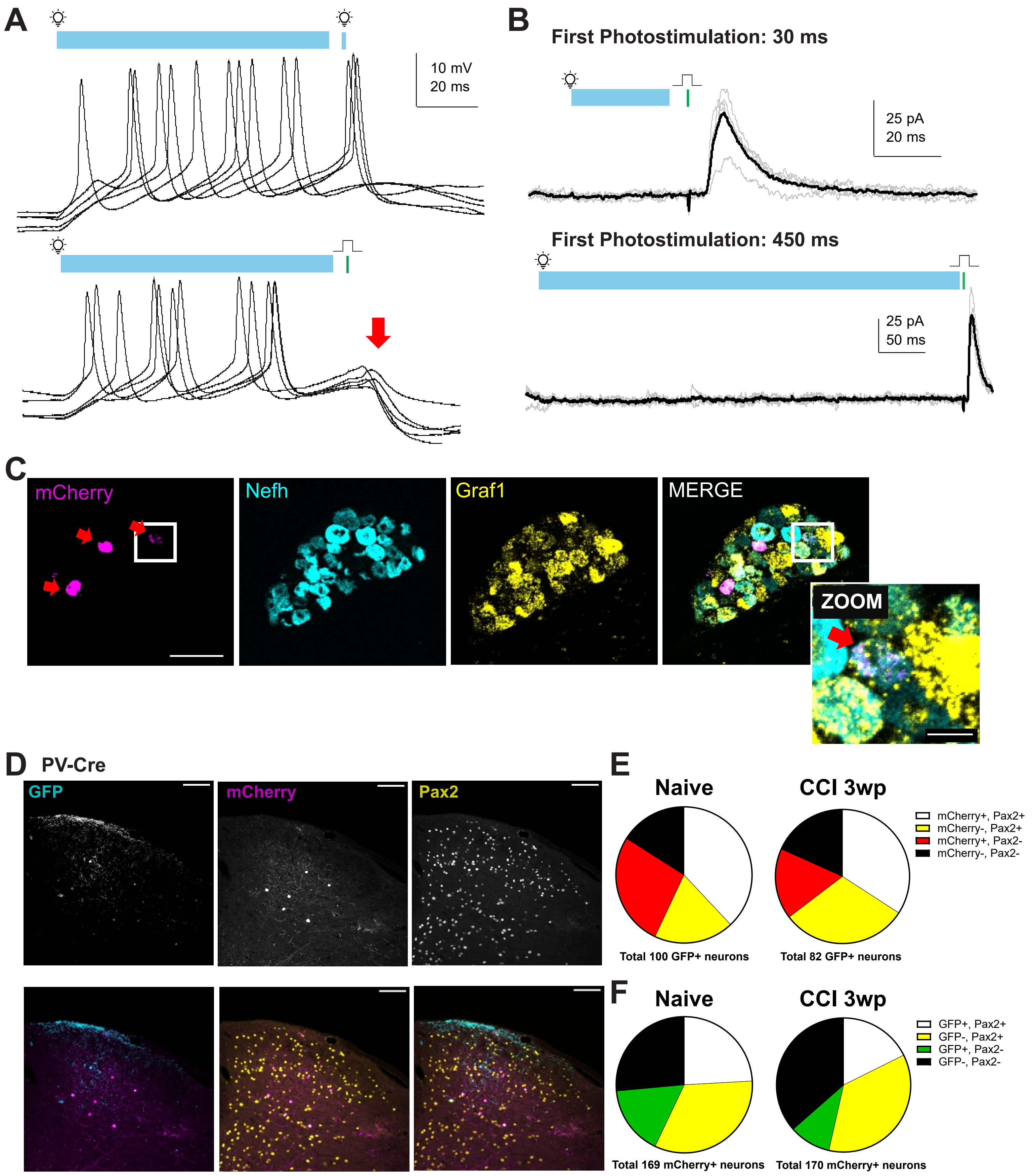

A
10 mV
20 ms
B
First Photostimulation: 30 ms
25 pA
20 ms
First Photostimulation: 450 ms
25 pA
50 ms
C
mCherry
Nefh
Graf1
MERGE
ZOOM
D
PV-Cre
GFP
mCherry
Pax2
E
Naive
CCI 3wp
mCherry+, Pax2+
mCherry-, Pax2+
mCherry+, Pax2-
mCherry-, Pax2-
Total 100 GFP+ neurons
Total 82 GFP+ neurons
F
Naive
CCI 3wp
GFP+, Pax2+
GFP-, Pax2+
GFP+, Pax2-
GFP-, Pax2-
Total 169 mCherry+ neurons
Total 170 mCherry+ neurons

### Figure S14

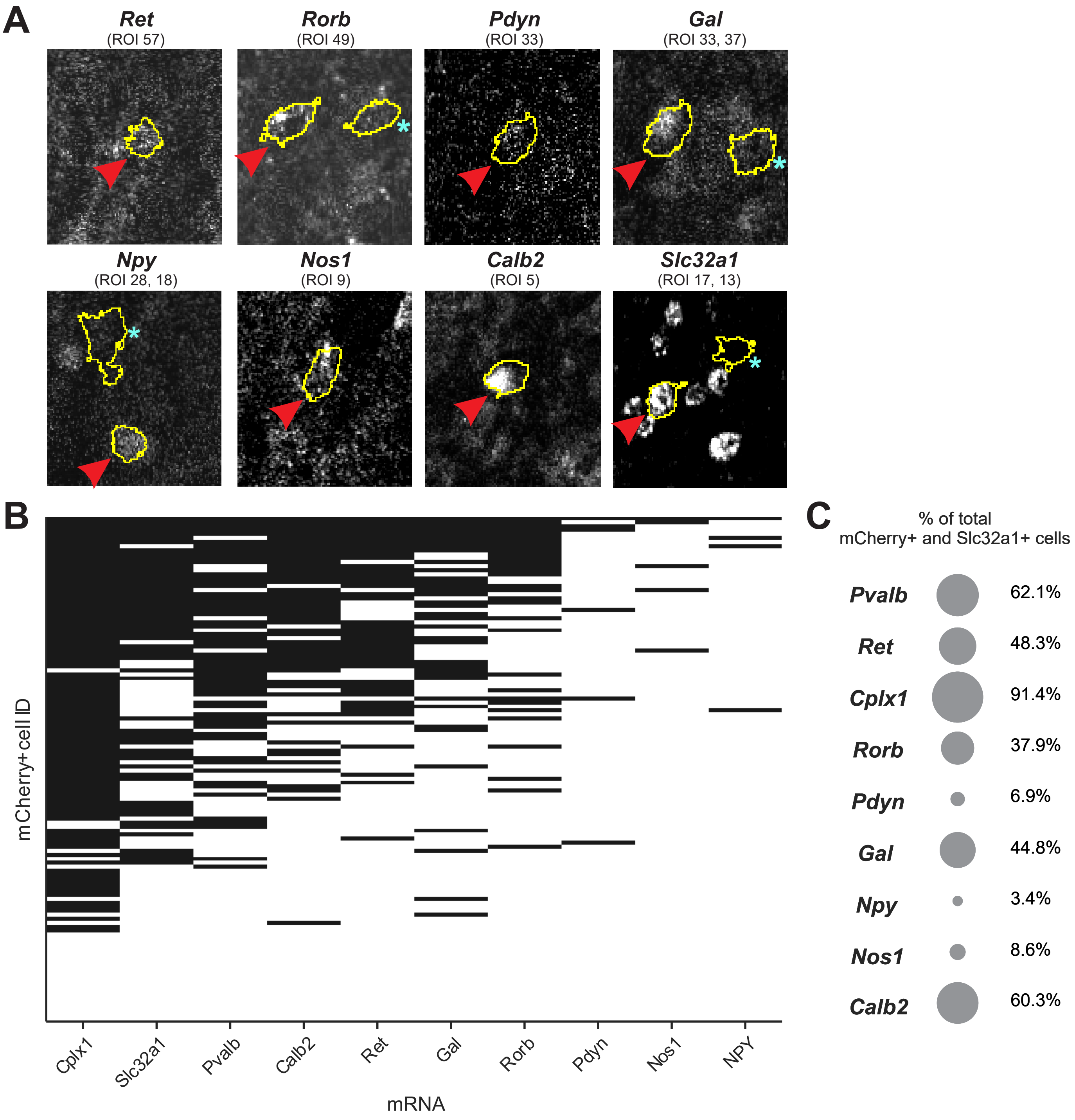
A
Ret
(ROI 57)
Rorb
(ROI 49)
Pdyn
(ROI 33)
Gal
(ROI 33, 37)
Npy
(ROI 28, 18)
Nos1
(ROI 9)
Calb2
(ROI 5)
Slc32a1
(ROI 17, 13)
B
mCherry+cell ID
Cplx1
Slc32a1
Pvalb
Calb2
Ret
Gal
Rorb
Pdyn
Nos1
NPY
mRNA
C
% of total
mCherry+ and Slc32a1+ cells
Pvalb 62.1%
Ret 48.3%
Cplx1 91.4%
Rorb 37.9%
Pdyn 6.9%
Gal 44.8%
Npy 3.4%
Nos1 8.6%
Calb2 60.3%

Figure S1 related to Figure 1: Characterization of primary afferent inputs onto PVNs

A-B. Representative traces of synaptically evoked responses in PVNs from naïve (A) and 3wp-CCI (B) mice. Monosynaptic Aβ inputs and monosynaptic or polysynaptic Aδ (red arrow) inputs were distinguished using varying stimulation intensities (25, 80, 100, and 500µA) and stimulation frequencies (0.1, 2, 4, 2, and 40Hz).

C. Proportion of PVNs receiving either exclusively Aβ inputs or combined Aβ+Aδ inputs in naïve (n=46 cells from 14 mice) and 3wp-CCI (n=29 cells from 11 mice) conditions. Chi-square contingency test.

D. Conduction velocities of Aβ and Aδ primary afferents synapsing onto PVNs in naïve and 3wp-CCI mice.

E-F. Quantification of evoked EPSC (eEPSC) latency (E) and the evoked current amplitude (F) of PVNs in naïve and 3wp-CCI mice. Unpaired t-test.

G-H. Representative paired-pulse responses evoked at interstimulus intervals of 500ms and 25ms (G), and the quantification of paired-pulse ratios (H) in naïve and 3wp-CCI conditions. Two-way ANOVA, Šídák's multiple comparison.

Figure S2 related to Figure 1: Recruiting different intensities of Aβ inputs onto PVNs

A. Representative voltage-clamp traces of synaptically evoked responses in PVNs elicited by increasing Aβ fiber stimulation intensities (5-75µA) in naïve, 5dp-CCI, and 3wp-CCI mice. Cells were held at -70mV.

B. Quantification of evoked current amplitudes across stimulations intensities (5µA-75µA), expressed as a percent of the amplitude evoked at 75µA electrical stimulation in naïve (n=5 cells from 2 mice), 5dp (n=5 cells from 2 mice), and 3wp-CCI (n=5 cells from 2 mice) conditions. Recordings were performed at -70mV to measure eEPSCs and at 0mV to measure eIPSCs, using stimulation frequencies of 2Hz and 40Hz.

C. Absolute evoked current amplitude at 75µA corresponding to the data in Panel B.

Figure S3 related to Figure 1: Additional UMAP clustering analyses

A-C. UMAP projections of synaptically evoked PVN responses from naïve, 5dp, and 3wp-CCI conditions, as shown in Figure 1, recolored for additional analyses. Responses are color-coded by sex (A; male, blue; female, red), by individual animal to assess batch effects (B; each mouse represented by a distinct color), and by intrinsic firing (C; color scale denotes the number of spikes evoked by a 150pA current injection).

D. Proportion of responses assigned to each cluster across naïve, 5dp, and 3wp-CCI conditions. One-way ANOVA, Tukey's multiple comparisons.

Figure S4 related to Figure 1: Cluster features after nerve injury

A-C. Z-scored synaptic response features across clusters and response classes in naïve, 5dp, and 3wp-CCI conditions. Shown are the average maximum amplitude (A), minimum amplitude (B), and area under the curve (AUC; C), grouped by cluster ID and classified as action potential (AP), excitatory postsynaptic potential (EPSP), or inhibitory postsynaptic potential (IPSP).

Figure S5 related to Figure 1: Changes in PVN mEPSCs after nerve injury

A. Representative voltage-clamp recordings of mini EPSCs (mEPSCs) from PVNs held at -70mV using a potassium-gluconate based internal solution and bath application of TTX (1µM), bicuculline (15µM), strychnine (500nM) in naïve, 5dp, and 3wp-CCI mice.

B. Cumulative probability distributions of mEPSC amplitudes from PVNs in naïve, 5dp, and 3wp-CCI mice. Pairwise two-sample Kolmogorov-Smirnov tests with Bonferroni correction.

C-H. Quantification of mEPSC in properties in PVNs from naïve, 5dp, and 3wp-CCI mice, including event frequency (C), decay time constant tau (D), half-width (E), rise time (F), decay time (G), and coefficient of variation of peak amplitude (H). One-way ANOVA, Tukey's multiple comparisons.

I. Representative images of a 3-dimensionally reconstructed PVcre;tdTomato neuron (red) and immunolabeling for Homer1 (white) and VGluT1 (green) in naïve and 3wp-CCI mice. Scale bar 30 µM.

J-L. Quantification of average Homer1 fluorescence intensity (J), surface area of PVcre;tdtomato neuron (K), and density of VGluT1/Homer1 appositions normalized to neuronal surface area (L) in naïve and 3wp-CCI mice. Unpaired t-test.

Figure S6 related to Figure 1: Changes in PVN mIPSCs after nerve injury

A. Representative voltage-clamp recordings of mini IPSCs (mIPSCs) from PVNs held at -70mV using a cesium chloride-based internal solution and bath application of TTX (1µM) and NBQX (10µM) in naïve, 5dp, and 3wp-CCI mice.

B. Cumulative probability distributions of mIPSC amplitudes from PVNs in naïve, 5dp, and 3wp-CCI mice. Pairwise two-sample Kolmogorov-Smirnov tests with Bonferroni correction. (*** $p < 0.001$ for naïve vs. 5dp; **** $p < 0.0001$ for 5dp vs. 3wp).

C-H. Quantification of mIPSC in properties in PVNs from naïve, 5dp, and 3wp-CCI mice, including event frequency (C), decay time constant tau (D), half-width (E), rise time (F), decay time (G), and coefficient of variation of peak amplitude (H). One-way ANOVA, Tukey's multiple comparisons.

I. Representative images of a 3-dimensionally reconstructed PVcre;tdTomato neuron (red) and immunolabeling for gephyrin (white) in naïve and 3wp-CCI mice. Scale bar 30 μM.

J-L. Quantification of average gephyrin fluorescence intensity in processes (J), somas (K), and surface area of PVcre;tdtomato neuron (L) in naïve and 3wp-CCI mice. Unpaired t-test.

Figure S7 related to Figure 2: Characterization of ChR2 activated primary afferent inputs onto PVNs

A. Representative current-clamp recordings of electrically evoked IPSPs (eIPSPs) in PVNs mediated exclusively by glycinergic transmission (top) or GABAergic transmission (bottom).

B. Quantification of eIPSP amplitudes in PVNs shown in panel A following sequential bath application of strychnine and then bicuculline + strychnine (top), or bicuculline and then bicuculline + strychnine (bottom).

C. Representative image of the lumbar spinal cord (left) and L4 dorsal root ganglion (DRG; right) of PVcre;tdTomato mice that received AAV injection to express ChR2-GFP restrict to peripheral primary afferents and terminals, with no detectable expression in spinal interneuron cell bodies in the dorsal horn. Scale bar 500 μM.

D. Quantification of PVN spike number after a 150pA step current injection of PVNs following TTX/4-AP bath application, confirming effective sodium channel blockade in naïve and 3wp-CCI mice.

E-F. Representative voltage-clamp recordings (E) and quantification (F) of optogenetically evoked IPSCs (oIPSCs; holding potential -40mV) and EPSCs (oEPSCs; holding potential -70mV) recorded from the same PVN following blue-light activation of peripheral afferents in the presence of TTX alone.

G. Heatmap of individual optogenetically evoked synaptic responses recorded from PVNs, ordered by increasing response amplitude, in naïve and 3wp-CCI conditions.

H. Quantification of maximum (top) and minimum amplitude (bottom) of optogenetically evoked synaptic responses in PVNs from naïve and 3wp-CCI conditions.

I-K. Current-clamp recordings and response classification from the same PVNs showing responses evoked by electrical stimulation and by optogenetic activation of primary afferents inputs in naïve (I), 5dp- (J), and 3wp-CCI (K) conditions.

Figure S8 related to Figure 3 and 4: Comparison of iPVN and ePVN intrinsic firing and primary afferent inputs

A. Whole-cell current-clamp representative traces of iPVN (top) and ePVN (bottom) response to step current injections of naive (Left) and 3wp-CCI (Right) mice.

B. Spike count of iPVN, ePVN, and combined PVN response to step current injections in naive and 3wp-CCI mice. iPVN: n = 28 cells from 8 naive mice and n = 29 cells from 7 CCI mice. ePVN: n= 30 cells from 8 naive mice and n = 26 cells from 6 CCI mice. *p-value=0.0472, Two-way ANOVA, Uncorrected Fisher's LSD compares combined PVN naïve vs 3wp-CCI.

C-D. Representative paired-pulse responses evoked at interstimulus intervals of 500ms and 25ms (C), and the quantification of paired-pulse ratios (D) in iPVN and ePVN, naïve and 3wp-CCI conditions. One-way ANOVA, Šídák's multiple comparison.

E-F. Quantification of evoked EPSC (eEPSC) latency (E) and the evoked current amplitude (F) of iPVNs and ePVNs in naïve and 3wp-CCI mice. One-way ANOVA, Brown-Forsythe and Welch's multiple comparison. *p-value=0.01

G. Success rate of action potential generation out of 100 Aβ stimulations (4Hz, 75µA, 25s) in iPVNs and ePVNs in naïve and 3wp-CCI mice. One-way ANOVA, Tukey's multiple comparison. **p-value = 0.0033, ****p-value <0.0001.

Figure S9 related to Figure 3 and 4: Additional UMAP clustering analyses for iPVN and ePVN

A-C. UMAP projections of synaptically evoked iPVN and ePVN responses from naïve, 3wp-CCI conditions, as shown in Figure 3 and 4, recolored for additional analyses. Responses are color-coded by sex (A; male, blue; female, red), by individual animal to assess batch effects (B; each mouse represented by a distinct color), and by intrinsic firing (C; color scale denotes the number of spikes evoked by a 150pA current injection).

D. Proportion of responses assigned to each cluster across iPVN and ePVN, naïve and 3wp-CCI conditions. One-way ANOVA, Tukey's multiple comparisons.

Figure S10 related to Figure 3 and 4: Cluster features after nerve injury for iPVN and ePVN

A-C. Z-scored synaptic response features across clusters and response classes in iPVN and ePVN, naïve and 3wp-CCI conditions. Shown are the average maximum amplitude (A), minimum amplitude (B), and area under the curve (AUC; C), grouped by cluster ID and classified as action potential (AP), excitatory postsynaptic potential (EPSP), or inhibitory postsynaptic potential (IPSP).

Figure S11 related to Figure 3 and 4: No changes to mEPSCs and sEPSCs of iPVNs and ePVNs after nerve injury

A-B. Quantification of mEPSC in properties in iPVNs (A) and ePVNs (B) from naïve and 3wp-CCI mice, including half-width, rise time, decay time, and coefficient of variation of peak amplitude. Unpaired t-test.

C-D. Quantification of sEPSC in properties in iPVNs (C) and ePVNs (D) from naïve and 3wp-CCI mice, including half-width, rise time, decay time, and coefficient of variation of peak amplitude. Unpaired t-test.

Figure S12 related to Figure 3 and 4: iPVNs and ePVNs are modality-specific to mechanical stimuli

A-B. Validation of the hM4D(Gi)-mCherry (A, images show mCherry expression) and HA-hM3D(Gq) (B, images show HA expression) virus in the spinal cord of GlyT2Cre;PVDre;Ai66D (left) and VGluT2Cre;PVDre;Ai66D (right) mice. Scale bar 500 μM.

C-D. Hargreaves' withdrawal latency (C) and acetone duration of nocifensive behaviour (D) in naïve GlyT2Cre;PVDre;Ai66D mice injected with hM4D(Gi)-mCherry. N= 10 mice saline, N=10 mice DCZ, paired t-test, n.s. not statistically significant.

E-F. Hargreaves' withdrawal latency (E) and acetone duration of nocifensive behaviour (F) in CCI 3wp GlyT2Cre;PVDre;Ai66D mice injected with HA-hM3D(Gq). N= 11 mice saline, N=11 mice DCZ, paired t-test, n.s. not statistically significant.

G-H. Hargreaves' withdrawal latency (G) and acetone duration of nocifensive behaviour (H) in naïve VGluT2Cre;PVDre;Ai66D mice injected with HA-hM3D(Gq). N= 9 mice saline, N=9 mice DCZ, paired t-test, n.s. not statistically significant.

E-F. Hargreaves' withdrawal latency (E) and acetone duration of nocifensive behaviour (F) in CCI 3wp VGluT2Cre;PVDre;Ai66D mice injected with hM4D(Gi)-mCherry. N= 10 mice saline, N=10 mice DCZ, paired t-test, n.s. not statistically significant.

Figure S13 related to Figure 5: Characterization of pre-synaptic spinal and primary afferents to PVNs through rabies tracing

A. Validation of PVcre;ChR2-tdTomato mice. Current clamp recordings show upon blue light activation, action potentials are generated; however, dorsal root electrical stimulation elicits IPSPs.

B. Different time intervals (30ms, 450ms) of the first photo-stimulation still cause the subsequent dorsal root electrical stimulation to elicit IPSCs.

C. Representative immunohistochemical images of dorsal horn section showing rabies-traced pre-synaptic neurons connected to PVNs. GFP-positive cells show source PVNs, while mCherry-positive cells show rabies-infected, transsynaptically traced neurons. Sections were immunolabelled for Pax2 to identify inhibitory interneurons. Scale bar 100 μM.

D-E. Quantification of rabies tracing results in naïve and 3wp-CCI mice. Shown are the proportions of total GFP-positive source neurons that are mCherry-positive or -negative and Pax2-positive or -negative (D), and the proportion of total mCherry-positive rabies-infected neurons that are GFP-positive or-negative and Pax2-positive or -negative (E). Scale bar 100 μM, 30 μM.

F. Fluorescent *in situ* hybridization images of L4 DRG showing rabies-traced presynaptic primary afferent neurons forming mono-synaptic connections onto PVNs. Rabies-labelled neurons are indicated by mCherry expression and are colabelled with Nefh and Grafl to indentify Aβ fibers. Red arrows show presynaptic labelled Aβ neurons. Scale bar 100 μM, 30 μM.

Naïve (n=2 mice): 12/18 mCherry+ cells were Nefh+/Graf1+. 6/18 mCherry+ cells were Nefh+/Graf1-. CCI (n=3 mice): 41/52 mCherry+ cells were Nefh1+/Graf1+. 11/52 mCherry+ cells were Nefh+/Graf1-.

<u>Figure S14 related to Figure 5: Identification of inhibitory neurons pre-synaptic to PVNs</u>

A. Representative images of HiPlex fluorescent *in situ* hybridization images of outlined rabies-traced mCherry+ cells that are positive for *Ret, Rorb, Pdyn, Gal, Npy, Nos1, Calb2,* and *Slc32a1*. Red arrows indicate cells with positive expression, cyan asterisks indicate cells with negative expression.

B. Quantification of panel A showing a binary heatmap of the expression of the different mRNA markers per cell. n=125 mCherry+ cells from 2 mice.

C. Quantification of the percentage of the colocalization of different neuron markers with the pre-synaptic mCherry+ and Slc32a1+ neurons of PVNs. n=58 mCherry+ and Slc32a1+ cells from 2 mice.

Supplementary Table 1: Action potential properties of the intrinsic firing of iPVNs and ePVNs in naïve and $CCI_{3wp}$. iPVN: n = 28 cells from 8 naive mice and n = 29 cells from 7 CCI mice. ePVN: n= 30 cells from 8 naive mice and n = 26 cells from 6 CCI mice.

| Mean ± SEM | iPVN Naïve (n=28) | iPVN CCI $_{3wp}$ (n =29) | ePVN Naïve (n=30) | ePVN CCI $_{3wp}$ (n=26) |
|---|---|---|---|---|
| Onset (mV) | -39.06 ± 1.87 | -44.41 ± 1.85 | -39.09 ± 1.24 | -40.55 ± 1.08 |
| Peak (mV) | -0.25 ± 1.67 | 5.88 ± 1.49 | 17.34 ± 1.39 | 16.30 ± 1.32 |
| Offset (mV) | -31.74 ± 1.50 | -42.48 ± 1.39 | -35.66 ± 1.37 | -38.36 ± 1.10 |
| Max Rise Slope (mV/ms) | 41.94 ± 4.12 | 50.40 ± 5.84 | 58.42 ± 6.24 | 60.57 ± 6.36 |
| Max Fall Slope (mV/ms) | -6.26 ± 11.34 | -22.74 ± 3.00 | -34.08 ± 3.99 | -34.93 ± 3.36 |
| Duration (ms) | 58.05 ± 17.03 | 54.85 ± 19.37 | 20.64 ± 5.27 | 22.36 ± 7.49 |
| Spike Threshold (mV) | -26.61 ± 1.18 | -32.58 ± 1.60 | -24.03 ± 0.81 | -24.32 ± 0.75 |
| Half Width (ms) | 2.83 ± 0.31 | 2.11 ± 0.30 | 2.08 ± 0.17 | 2.00 ± 0.17 |
| Base Width (ms) | 5.94 ± 1.01 | 3.78 ± 0.34 | 4.13 ± 0.31 | 3.59 ± 0.28 |
| Rise Time (ms) | 2.02 ± 0.22 | 1.52 ± 0.13 | 1.86 ± 0.14 | 1.65 ± 0.13 |
| Fall Time (ms) | 5.96 ± 0.86 | 7.36 ± 4.99 | 2.25 ± 0.17 | 1.97 ± 0.15 |
| Rise Rate (mV/ms) | 20.33 ± 1.61 | 31.44 ± 3.34 | 28.86 ± 2.78 | 31.49 ± 3.05 |
| Fall Rate (mV/ms) | -16.37 ± 2.01 | -19.98 ± 3.21 | -24.88 ± 3.06 | -25.24 ± 2.96 |
